# Finding the relatives of a critically endangered island endemic reveals a heterogeneous evolution of land snail mitogenomes (Gastropoda: Helicidae)

**DOI:** 10.1101/2025.03.24.642578

**Authors:** Ondřej Korábek, Katerina Vardinoyannis, Aydin Örstan, Nikolaos Psonis

## Abstract

We analysed the phylogenetic relationships of *Aristena rechingeri*, an extremely rare, large, flat-shelled snail from the Aegean island of Karpathos. Low-depth Illumina sequencing data obtained from an empty shell were available, but there was no reference genome or sequences of nuclear loci from the related taxa, so we attempted to resolve the phylogeny with mitogenome sequences. We confirmed that *Aristena* is sister to the globular-shelled genera *Helix* and *Maltzanella* and not to the similarly flat-shelled and geographically proximate *Isaurica* from Anatolia. Concurrently, we clarified the status of two other flat-shelled taxa (*Isaurica callirhoe*, *Levantina menkhorsti*). However, even data from complete mitogenomes do not resolve most of the intergeneric relationships in the tree. The mitogenome sequences show variation in nucleotide compositions and substitution rates among species, making phylogenetic inference difficult even with complex mixture models. In general, our examination of the properties of the mitogenomes point to their dynamic and varied evolution, which misleads phylogenetic analyses even at shallow phylogenetic depths. Furthermore, we found that duplications within the mitogenomes are not very rare and that protein-coding genes followed by tRNAs typically end with an incomplete stop codon. The latter presents an issue for automatic annotation tools.

## INTRODUCTION

The western Palearctic family Helicidae Rafinesque, 1815 is the best studied group of land snails. It contains several widespread, large, and edible species, including the brown garden snail *Cornu aspersum* (Müller, 1774) that is important in heliciculture and has become invasive in many parts of the world (Guiller et al., 2012), the grove snail *Cepaea nemoralis* (Linnaeus, 1758) that has been a prominent model for evolutionary studies (e.g. Cain & Sheppard, 1954; Cook, 1998; Jones et al., 1977; Lamotte, 1959; Silvertown et al., 2011), and the Roman snail *Helix pomatia* Linnaeus, 1758, a well studied species that is a characteristic snail of Central Europe (e.g., Meisenheimer, 1912; Korábek et al., 2018). Not surprisingly, one of these helicids (*C. nemoralis*) was among the first land snails with complete mitogenome sequenced (Terrett et al., 1996).

Mitochondrial sequence data have been immensely important in the systematics and biogeography of land snails, and Helicidae are no exception (e.g. Fiorentino et al., 2010; Greve et al., 2010; Chueca et al., 2015; Psonis et al., 2015; Walther et al., 2016; Holyoak et al., 2020; Neiber, et al., 2021; Neiber et al., 2022; Korábek, Juřičková, et al., 2022; Korábek et al., 2023). Despite advances in methods of sequencing and analysis of genome-wide data and well-known disadvantages of the mitochondrial data, mitochondrial marker sequences remains an effective tool for the exploration of diversity in a situation when many land snail genera still lack any sequence data, new species keep being described without even a DNA barcode, and most of the earlier data are mitochondrial sequences. Hence, they continue to be used a lot (e.g. Bullis & Rundell, 2024; Hyman & Köhler, 2024; Nekola et al., 2024; Saito et al., 2024; Stanisic et al., 2024). In fact, sometimes there is no other option than to resort to mitochondrial data. It may be impossible to perform additional sequencing and genomic analyses because the populations in question live in presently inaccessible areas, are difficult to collect, or have become extinct. Mitochondrial sequences from earlier studies or obtained from poorly preserved samples may be the only data available for study. However, even after all the years of research relying on mitochondrial sequences, the knowledge of the basic characteristics of the mitogenome evolution in land snails is poor. That seriously hampers our ability to assess the reliability and significance of phylogenetic results obtained from analyses of mitochondrial sequences.

An early study of the mitochondrial variation in *C. nemoralis* revealed deep intraspecific divergences (Thomaz et al., 1996) and one of the proposed possible explanations was an exceptionally high substitution rate in that species. A recent study of the same species found substantial levels of heteroplasmy involving surprisingly high divergences and variation in the number of tRNA gene copies (Davison et al., 2024), suggesting a fast and dynamic evolution of mitogenome sequences in this helicid. Indeed, the substitution rate of *C. nemoralis* appears to stand out as particularly high among Helicidae (see figure S1 in Korábek, Juřičková, et al., 2022). However, there is not much data to compare with. As of January 2025, there were only six helicid genera with published mitogenome sequences (Groenenberg et al., 2012; Wang et al., 2018; The Darwin Tree of Life Project Consortium et al., 2022; Davison et al., 2024; Korábek & Hausdorf, 2024) out of 52, and only in *Helix* Linnaeus, 1758 and *Cepaea* Held, 1838 more than one species was analysed (Davison et al., 2024; Korábek & Hausdorf, 2024). It is therefore unclear how much the substitution rates vary between and within genera, how much is this variation consistent along the mitogenome sequences, what other parameters vary (e.g. nucleotide composition, strength of selection), and, most importantly, how does all this impact studies using mitochondrial data to gain insights into phylogeny and phylogeography. A new study on *Helix* (Korábek & Hausdorf, in prep.) suggests that the impact is significant and negatively affects phylogenetic analyses even between very closely related species.

In this study, we aimed to resolve the phylogenetic position of the monotypic genus *Aristena* Psonis, Vardinoyannis & Poulakakis, 2022, endemic to the Aegean island of Karpathos, using mitogenomic data. Short read sequences of *Aristena rechingeri* (Fuchs & Käufel, 1936) were obtained by Psonis et al. (2022) from an empty shell, because despite repeated efforts since its description (Pfeiffer & Wächtler, 1939; Glaubrecht, 1994; Grano & Cattaneo, 2021; Psonis et al., 2022) the species was for long not found alive. This changed only after the sequencing (Porfyris, 2023), and the exact locality where living specimens were found was not disclosed.

*Aristena* is one of the cases where mitochondrial data remain as yet not easily replaceable. Given the shallow shotgun sequencing, only a part of *Aristena*’s mitogenome was previously assembled together with histones H3 and H4 and the nuclear rDNA gene cluster (Psonis et al., 2022). Due to lack of comparable data from other species, the phylogenetic position of *Aristena* was previously inferred only on the basis of partial *cox1* and *rrnL* genes (mitochondrial) and the internal transcribed spacer 2 (ITS2; nuclear) (Psonis et al., 2022). There is no reference genome from a closely related species to map the Illumina reads to, nor are there published sequences of a sufficient number of nuclear loci. We thus decided to use mitogenome sequences to re-assess the relationships of Aristena, as these were possible to obtain from related genera easily. Using an approach modified from the original study we were able to assemble here a nearly complete mitogenome of *A. rechingeri* from the same raw reads produced by Psonis et al. (2022). In order to be able to resolve the phylogenetic position of the genus, we also assembled eight additional complete or nearly complete mitogenomes from related genera and combined these data with mitogenomic sequences from *Helix* Linnaeus, 1758, *Maltzanella* Hesse, 1917, and four outgroups.

The previous phylogenetic analysis (Psonis et al., 2022) suggested a close relationship to *Helix* and its sister group *Maltzanella*, but were statistically inconclusive (support from a Bayesian, but not a Maximum Likelihood analysis). The anatomy of the genital system, which was the principal source of characters for the genus-level taxonomy of the rock-dwelling helicids before molecular phylogenetic studies, is unknown for *Aristena*. Geographic proximity, which is often a good predictor of phylogenetic relationships in land snails, would suggest a close relationship to *Isaurica* Kobelt, 1901, which has been indeed proposed before (Pfeiffer & Wächtler, 1939). *Isaurica* includes five currently accepted species, but only three of them have been examined anatomically (Subai, 1994) and sequence data have been published for only two species (Korábek, Juřičková, et al., 2022). One species, *Isaurica callirhoe* (Rolle, 1894), is particularly problematic. It was long known only from the type series collected at a very imprecisely described locality (Rolle & Kobelt, 1896); the single later report of the species being found (Ceylan et al., 2008) went largely unnoticed. Nordsieck (2017) speculated that *I. callirhoe* should, in fact, be placed within *Assyriella* Hesse, 1909 (a junior synonym of *Levantina* Kobelt, 1871), but that was before all the western occurrences of *Levantina*/*Assyriella* were reinterpreted as anthropogenic (*L. spiriplana* (Olivier, 1801)) or as referring to a different genus (*A. rechingeri*) (Korábek, Glaubrecht, et al., 2022; Psonis et al., 2022). Because this species remains enigmatic and thus could not be excluded as a candidate for a close relationship to *Aristena*, we also revised its status, although only with the sequences of short mitochondrial markers.

The mitogenomic dataset we collated allowed us to provide a detailed analysis of the structural properties of the mitogenomes and to examine the variability in mitogenome sequence characteristics among helicids. We focused on the variation in gene order, start and stop codons, substitution rates, and nucleotide composition. Our results are relevant for the use of mitochondrial data in land snail phylogenetics, demonstrating that the high heterogeneity in the substitution process among lineages influences phylogenetic analyses. We also highlight some limitations of automatic annotation tools when used for the annotation of land snail mitogenomes.

## MATERIALS AND METHODS

### Mitogenomes: sequencing, assembly, annotation

#### Selection of taxa

All the studied species (Table 1) belong to the tribe Helicini, a group of 10 well-supported monophyletic genera with unresolved relationships among them (Korábek, Juřičková, et al., 2022; Neiber et al., 2022). We combined previously assembled mitogenome sequences from *Helix* and *Maltzanella* (Korábek & Hausdorf, 2024; Korábek & Hausdorf, in prep.) with additional RNA or DNA short read sequencing data of representatives of the tribe. For both *Levantina* and *Caucasotachea* Boettger, 1909, in which the oldest intrageneric splits are deep, we added a second species to break up their branches. As outgroups, we used species from other tribes of the subfamily Helicinae, for which mitogenomes were available at the NCBI GenBank database (Sayers et al., 2024) as of January 2025: *Cornu aspersum* (The Darwin Tree of Life Project Consortium et al., 2022), *Theba pisana* (Müller, 1774) (Wang et al., 2018), *Cepaea nemoralis*, *Cepaea hortensis* (Müller, 1774) (The Darwin Tree of Life Project Consortium et al., 2022; Davison et al., 2024).

**Table 1.** List of species used for molecular analyses with locality information. Further details (GPS coordinates, voucher data, NCBI GenBank and SRA accession numbers) are available from the Supplementary Table S1.

| ID | species | locality | isolate (lab code) | reference |
| --- | --- | --- | --- | --- |
| HE001286 | *Amanica praezellens* (Nägele, 1901) | Turkey: Hatay Prov.: Yeniköy 4 km above Payas | IS5 | this study |
| HE003148 | *Aristena rechingeri* (Fuchs & Käufel, 1936) | Greece: Karpathos Isl., NW of Myrtonas | ADNA1001251 | Psonis et al., 2022, this study |
| HE003272 | *Caucasotachea atrolabiata* (Krynicky, 1833) | Georgia: Imereti: Tskaltubo, Kumistavi, entrance of Prometheus Cave | 159145 | this study |
| HE000991 | *Caucasotachea vindobonensis* (Pfeiffer, 1828) | Russia: Moscow Region: Lobnya | VIND | Korábek & Hausdorf, 2024 |
| HE003253 | *Codringtonia parnassia* (Roth, 1855) | Greece: Central Greece: 0.5 km from Analipsi on road to Kyriaki | 42947f | this study |
| HE002843 | *Helix albens* Rossmässler, 1839 | Russia: Krasnodar: Verkhnebakanskiy, 2 rkm SW of: pass between Anapa and Novorossiysk | 6986 | Korábek & Hausdorf, in prep. |
| HE001338 | *Helix buchii* Dubois de Montpéroux, 1840 | Turkey: Trabzon Prov.: Sümela monastery | BC2 | Korábek & Hausdorf, in prep. |
| HE002865 | *Helix buchii* Dubois de Montpéroux, 1840 | Georgia: Mtskheta-Mtianeti: T'ianet'i: 4.5 rkm towards Tbilisi | 7000 | Korábek & Hausdorf, in prep. |
| HE003275 | *Helix cincta* Müller, 1774 | Italy: Veneto: Verona: Belluno Veronese, cemetery | 141579a | Korábek & Hausdorf, 2024 |
| HE003273 | *Helix figulina* Rossmässler, 1839 | Greece: Central Greece: Chaironeia to Thouria, limestone hill by the road | 42946 | Korábek & Hausdorf, 2024 |
| HE003268 | *Helix godetiana* Kobelt, 1878 | Greece: Dodecanese: Astypalaia Isl., Koutsomytis islet | 6519 | Korábek & Hausdorf, 2024 |
| HE003276 | *Helix ligata* Müller, 1774 | Italy: Lazio: Rieti: Amatrice, gas station on road SS 4 near Bagnolo | 141559a | Korábek & Hausdorf, 2024 |
| HE002405 | *Helix lucorum* Linnaeus, 1758 | Serbia: Pomoravlje Dist.: highway north of Čuprija | LUC | Korábek & Hausdorf, 2024 |
| HE003257 | *Helix lutescens* Rossmässler, 1837 | Slovakia: Košický kraj: Kašša | 42983f | Korábek & Hausdorf, 2024 |
| HE002964 | *Helix nucula* Mousson, 1854 | Lebanon: Beqaa Gov.: N of Aammiq | 7057 | Korábek & Hausdorf, in prep. |
| HE003132 | *Helix pathetica* Mousson, 1854 | Turkey: Erzincan Prov.: road D100 between Çavuşköy and Kalıntaş | 7582 | Korábek & Hausdorf, in prep. |
| HE003251 | *Helix pelagonesica* (Rolle, 1898) | Greece: Central Macedonia: SE of Nikiti | 42932f | Korábek & Hausdorf, 2024 |
| HE002600 | *Helix philibinensis* Rossmässler, 1839 | Bulgaria: Blagoevgrad Prov.: Kresna Gorge S of Cherniche, old road | 6271 | Korábek & Hausdorf, in prep. |
| HE002644 | *Helix pomacella* Mousson, 1854 | Bulgaria: Yambol Prov.: N of Granitovo, by road to Elhovo | 6598 | Korábek & Hausdorf, in prep. |
| HE000505 | *Helix pomatia* Linnaeus, 1758 | Germany: Bayern: Unterdietfurt, Huldessen | P_488 | Korábek et al., 2019 |
| HE003277 | *Helix pomatia* Linnaeus, 1758 | Czechia: Jihomoravský kraj: Pouzdřany, NPP Pouzdřanská step – Kolby | 141580a | Korábek & Hausdorf, 2024 |
| HE002954 | *Helix pronuba* Westerlund, 1879 | Greece: Crete: Elafonisos, rocky slope | 7055 | Korábek & Hausdorf, in prep. |
| HE001273 | *Helix salomonica* Nägele, 1899 | Turkey: Hakkâri Prov.: Hakkâri, castle | SL1 | Korábek & Hausdorf, in prep. |
| HE000808 | *Helix secernenda* Rossmässler, 1847 | Montenegro: Lovćen Mts. | SEC | Korábek & Hausdorf, 2024 |
| HE002946 | *Helix thessalica* Boettger, 1886 | Czechia: Jihomoravský kraj: Mohelno, left bank of Jihlava | 7315 | Korábek & Hausdorf, 2024 |
| HE003274 | *Helix thessalica* Boettger, 1886 | Romania: Buzău County: SW of Ciuta | 141581a | Korábek & Hausdorf, 2024 |
| HE001725 | *Helix vladika* (Kobelt, 1898) | Montenegro: between Miloševići and Krnovo | VLAD | Korábek & Hausdorf, 2024 |
| HE001246 | *Isaurica schuetti* Subai, 1994 | Turkey: Antalya Prov.: Turunçova env., near Limyra | IS41 | this study |
| HE002250 | *Levantina escheriana* (Bourguignat, 1864) | Turkey: Mardin Prov.: Elmahahçe | CGR22912 | this study |
| HE002487 | *Levantina menkhorsti* (Gümüş & Neubert, 2012) | Turkey: Bitlis Prov.: Küçüksu köyü, 11 km E of Tatvan, limestone rocks on the left side of the road to Van | LV38 | this study |
| HE002944 | *Lindholmia nordmanni* (Mousson, 1854) | Turkey: Artvin: Su Kavuşumu: 1 rkm SE from crossroads Erzurum/Yusufeli/Artvin towards Yusufeli | 6930 | this study |
| HE001230 | *Maltzanella dickhauti* (Kobelt, 1903) | Turkey: Afyonkarahisar Prov.: Burunkaya near Dinar | MT7 | Korábek & Hausdorf, in prep. |
| HE003254 | *Neocrassa neocrassa* (Zilch, 1952) | Albania: Gjirokastrë County: Jorgucat, shallow valley immediately N of | 42963f | this study |
| HE003242 | *Isaurica callirhoe* (Rolle, 1894) | Turkey: Antalya Prov.: Akdağ mountain, steep limestone slope | 8049 | this study |
| HE002240 | *Isaurica lycia* (von Martens, 1889) | Turkey: Muğla Prov.: Kumlubük S of Marmaris | CGR22915 | Korábek et al., 2022 |
| HE002426 | *Isaurica lycia* (von Martens, 1889) | Turkey: Antalya Prov.: Demre, N of the ancient town of Myra, dirt road ending at a single building at the NE facing slope | ZMH3166 | Neiber et al., 2022 |
| HE002568 | *Isaurica lycia* (von Martens, 1889) | Turkey: Antalya Prov.: Demre, Kale | 6089 | this study |
| HE003236 | *Isaurica lycia* (von Martens, 1889) | Greece: Dodecanese: Megisti Isl. (Kastellorizo Isl.), Papalazarou fields | 7723 | this study |
| HE003237 | *Isaurica lycia* (von Martens, 1889) | Greece: Dodecanese: Megisti Isl. (Kastellorizo Isl.), Vigla mt., around plateau | 7722 | this study |
| HE002569 | *Isaurica pamphylica* Subai, 1994 | Turkey: Antalya Prov.: Altınkaya, ruins of Selge | 6088 | this study |
| HE002529 | *Isaurica schuetti* Subai, 1994 | Turkey: Antalya Prov.: 1 km S of junction to Olympos, 12 road km from Phaselis | IS10 | Neiber et al., 2022 |
| outgroup | *Cepaea nemoralis* (Linnaeus, 1758) | laboratory stock | n/a | Davison et al., 2024 |
| outgroup | *Cepaea hortensis* (Müller, 1774) | United Kingdom: Northamptonshire: Kettering, Geddington, 2 Queen Eleanor Road | n/a | DTOL, 2024 |
| outgroup | *Theba pisana* (Müller, 1774) | Spain: Mallorca: Can Pastilla | n/a | Wang et al., 2018 |
| outgroup | *Cornu aspersum* (Müller, 1774) | United Kingdom: Bedfordshire: Luton, Stopsley, Hollybush Road | n/a | DTOL, 2024 |

The complete list of samples along with accession numbers for the data deposited in NCBI’s nucleotide and SRA databases is provided in Supplementary Table S1.

#### Sequencing and assembly

The Helicini mitogenomes included in the analyses were sequenced and assembled on different occasions within a period of 10 years, so the methods differed among species. In brief, the complete or nearly complete mitogenomes were assembled either from RNA-seq reads with Trinity 2.11.0 (Grabherr et al., 2011) or from whole genome shotgun sequencing reads with MITObim 1.9.1 (Hahn et al., 2013). In both cases the sequencing was performed on Illumina instruments (different platforms). Assembly of the mitogenome of *A. rechingeri* is detailed below, details of the methods used for the other species are provided in the Supplementary Appendix S1.

To assemble the mitogenome of *A. rechingeri*, we used the Illumina reads obtained by (Psonis et al., 2022) from an empty shell as available at the NCBI SRA database (runs SRR14773380 and SRR14773381; Table S1) and concatenated the two runs. Quality and adapter filtering was done with TrimGalore `(--length 30 -q 20 --stringency 1 -e 0.1`).

The low mitogenome coverage of the reads (∼12×) and the short length of single-end reads did not allow for *de novo* assembly. Furthermore, the metagenomic character of the data, which also contains various microbial or other environmental DNA reads, calls for caution when assembling the sequence. Psonis et al. (2022) used a cautious assembly approach that allowed for high confidence in the assembly, but this resulted in not being able to recover large parts of the mitogenome. The principal limitation of the assembly approach is in the initial baiting step used by MITObim 1.9.1 (Hahn et al., 2013), where an exact match to a reference sequence is required. The length of the match previously used (21 bp) was a cautious choice, but too restrictive for the relatively distantly related reference sequences available. In protein coding genes, a match of seven codons in a row with another genus is expected to be rare due to high variability at least in the third codon positions. However, despite lack of an exact match the reads from *Aristena* are still expected to be similar in sequence to the reference, so it should be possible to find these by a method that evaluates an overall sequence similarity over the whole read length. Reads identified in this way can then be used to construct the starting reference sequence for MITObim. In addition, the read pool can be filtered for exogenous and other useless reads that may interfere with the assembly or be wrongly incorporated. In this way, we aimed at higher sensitivity while maintaining specificity.

To obtain the reference sequence for the first baiting step of MITObim, we identified reads that can be identified as Stylommatophoran mitochondrial sequences and exhibit clear sequence homology to the mitogenome of *Helix pomatia*. We used Kraken2 2.1.2 (Wood et al., 2019) to find reads classifiable as from a Stylommatophoran mitochondria. We downloaded mitochondrial sequences from the NCBI nucleotide database (on 30th January 2023) using nsdpy 0.2.3 (Hebert & Meglécz, 2022) (`-v -r –(eukaryota[organism] AND isauricamitochondrion[filter])”`) and constructed from them a Kraken2 database (`-- kmer-len 30 --minimizer-len 26 --minimizer-spaces 6`). The reads were then classified using Kraken2 and this database and only the reads being classified to Stylommatophora were extracted with extract_kraken_reads.py from KrakenTools 1.2 (Lu et al., 2022). This yielded 2,282 reads, of which 52 were classified as exogenous DNA when run against a pre-made database of archaeal, bacterial, viral, plasmid, human, protozoan, and fungal sequences (k2_pluspf_16gb_20221209; https://benlangmead.github.io/aws-indexes/k2) and therefore were removed. The remaining reads were then used to build a BLAST database (Camacho et al., 2009). We used blastn and the *H. pomatia* mitogenome as a query to search against the database with a word size set to 7, keeping reads with bitscore >40 (394 reads). The retrieved hits were aligned to the *H. pomatia* mitogenome with MAFFT 7.520 (Katoh & Standley, 2013) (`--addfragments --auto`) and their consensus sequence was called with UGENE 40.0 (Okonechnikov et al., 2012). This consensus was then used to make a chimaeric reference sequence from the *H. pomatia* mitogenome sequence. The chimaeric sequence served as a reference for a MITObim run on a set of the 2,230 reads identified above as Stylommatophoran (`--kbait 21`). The resulting gapped assembly was then used as reference for further MITObim runs.

To save some time and reduce the risk of incorporating exogenous reads, we filtered the original read pool in two ways before running MITObim again. First, we used PRINSEQ lite 0.20.4 (Schmieder & Edwards, 2011) to further remove low-complexity reads (e.g. reads with tandem repeats). Then we ran Kraken2 using the k2_pluspf_16gb_20221209 database and removed all classified reads as possible of exogenous origin. The filtered read pool and the gapped reference made in the previous step was then used for a MITObim assembly with `-- kbait 21`. This produced an assembly with only two gaps: one short in *rrnL* and one longer in the *nd1* gene. The latter gap was closed when we set `--kbait 18` while the assembled sequence remained otherwise identical.

There was a short deletion in the *rrnS* gene compared to the assembled sequence of Psonis et al. (2022). However, the presumably missing part of the sequence was the only place where the consensus of aligned reads from the blastn search differed significantly from the assembly, suggesting problems already at the stage of reference sequence construction. A similar gap appeared within *rrnL* if the reference sequence was not the chimaera of the reads found by BLAST and *H. pomatia*, but only the reads separated by Ns. We therefore repeated the assembly with `--kbait 21` using as reference *rrnS* sequences of species representing different genera: *Helix pomatia*, *Isaurica schuetti*, *Codringtonia elisabethae* Subai, 2005, *Neocrassa neocrassa*, *Amanica praecellens*, and *Levantina spiriplana*. All these assemblies resolved the gap as in Psonis et al. (2022), but the *L. spiriplana* reference introduced a 13 bp insert (creating a tandem repeat of this sequence) elsewhere within the *rrnS* gene. Finally, we used BWA 0.7.17 (Li & Durbin, 2009; Oliva et al., 2021) to map reads back onto the assembled sequence (`bwa aln -l 1024`), calculated the coverage and checked it for unexpected drops or peaks.

To compare the levels of exogenous DNA between the *A*. *rechingeri* (from shell), *I*. *schuetti* (dried, partially decomposed tissue), and other samples used for WGS (tissues fixed in ethanol), we run Kraken2 with the k2_pluspf_16gb_20240605 database.

### Annotation

Because of the frequent use of several alternative start codons and incomplete stop codons (Ghiselli et al., 2021), which is however not reflected in existing annotations, it is not possible to only use automatic annotation tools or most of the published annotations of land snail mitogenomes as these incorrectly identify the limits of the protein coding genes. We located the ends of the genes followed by tRNA genes by searching for reads from RNA-seq that extend into the poly(A) tail in several species. This includes not only protein-coding genes, but also *rrnL* where the transcript is also polyadenylated. Especially in samples sequenced in a single-end setting the tails were in many cases assembled by Trinity. The start codons were identified by aligning the genes across species and searching for a conserved position of potential start codon.

To annotate tRNAs, we accepted the annotation produced by MITOS2 (Donath et al., 2019), with a few exceptions and manual adjustments. The exact start and end positions should not be taken as definitive. *trnS2* was rarely located by MITOS2 (it has an aberrant structure; Ozerova et al., 2024) and a few others (e.g. *trnR* and *trnI*) were sometimes also not found. The tRNAs not located by MITOS2 were annotated by MITOS (Bernt et al., 2013). In a few cases, a one base overlap between tRNA and protein-coding genes was proposed by the software (e.g. *cytb* and *trnD*, *cox2* and *trnY*, *atp8* and *trnN*, *atp6*, and *trnR*). In those cases the overlapping base was excluded from the tRNA range. The start and end positions indicated by MITOS2 were in many cases not congruent between species. In particular, MITOS2 suggested in some species a large overlap between *trnK* and *cox1*. The limits of *trnK*, whose sequence is relatively conserved, thus needed in those cases a manual adjustment to remove the overlap and make the annotation consistent between species.

### Sanger marker sequences of *Isaurica*

To clarify the phylogenetic position of *I. callirhoe* and exclude the possibility that it was related to *A. rechingeri*, we obtained sequences of the *rrnL*, *rrnS*, and *cox1* genes from *I. callirhoe* (individual stored in the Carnegie Museum of Natural History, Pittsburgh, U.S.A., CM188835) and additional individuals of *Isaurica*. DNA was extracted by a modified Sokolov’s method (Sokolov, 2000; Scheel & Hausdorf, 2012). PCR was performed as described in Korábek, Juřičková, et al. (2022), targeting the regions delimited by the primer pairs 16Scs1+16Scs2 (Chiba, 1999), 12SGast_fwd2+12SGast_rev3 (Cadahía et al., 2014), and LCO1490+HCO2198 (Folmer et al., 1994), as modified by Hausdorf et al. (2003).

### Phylogenetic analyses

#### Intra-generic phylogeny of Isaurica

The *cox1*, *rrnL*, and *rrnS* sequences were aligned with MAFFT and concatenated. Maximum Likelihood was performed with IQ-TREE 2.3.6 (Kalyaanamoorthy et al., 2017; Minh et al., 2020) under an automatically selected mixture substitution model (Ren et al., 2024). Branch support was assessed with 500 bootstraps.

#### Relationships among genera

A study on *Helix* (Korábek & Hausdorf, in prep.) concluded that even with complete mitochondrial sequences, reconstructing the mitochondrial phylogeny is difficult due to variation in substitution rates and nucleotide composition among branches. The relationships between the genera that are of interest here are even more challenging: the divergences are older and the branches are longer without the possibility of breaking them by addition of more species. We therefore tried four different approaches to see if that changes the topology and support of the phylogeny recovered. We evaluated how well each method dealt with the among-branch heterogeneity by looking at the topology of three clades that are known to be difficult to resolve and that contain species with significantly different branch lengths. Two such cases are within the genus *Helix* and one in the outgroup.

1. *Helix nucula* should be sister to *H. salomonica*, not to *H. figulina*. *Helix figulina* has the shortest branch of the three and *H. salomonica*, with the longest branch, may incorrectly appear as the basal branch (Korábek & Hausdorf, in prep.).
2. *Helix ligata* should be sister to *H. pronuba*, not to *H. cincta*. *Helix cincta* has the shortest branch of the three, while the *H. pronuba* branch is distinctly long and also has a different base composition (Korábek & Hausdorf, in prep.).
3. In the outgroup, *Theba* should be sister to *Cornu* and not to *Cepaea* (Korábek, Juřičková, et al., 2022; Neiber et al., 2022). Both *Theba* and *Cepaea* are long-branch taxa, so their grouping would suggest that the method does not entirely overcome long branch attraction (Kapli et al., 2021).

We first performed partitioned Maximum Likelihood analyses of both the nucleotide and amino-acid alignments, which were partitioned by gene and, in case of the nucleotide alignment, by codon position. Best fit models were selected with ModelFinder (Kalyaanamoorthy et al., 2017) in IQ-TREE (Minh et al., 2020), allowing the merging of the initial partitions. The search for the best tree was repeated 10 times and the statistical support values from 200 bootstraps were then mapped to that tree. In the study including *Helix* only (Korábek & Hausdorf, in prep.), it appeared that applying a mixture model and alignment filtering can help to overcome phylogenetic artefacts caused by uneven substitution rates and compositions among taxa when analysing the nucleotide data. Thus, we filtered the nucleotide alignment with ALISCORE 2.2 (Kück et al., 2010) and removed the third codon positions that are the fastest-evolving sites suffering from the greatest compositional heterogeneity (Table 2). ALISCORE was run on the amino-acid alignment with the tree from the partitioned analysis of nucleotide alignment and whole codons were removed with Alicut 2.31 (Kück, 2017). Then, we run the analysis using a mixture model selected by MixtureFinder (Ren et al., 2025). Finally, we used PhyloBayes MPI 1.7 (Lartillot et al., 2013) to perform a Bayesian inference on the filtered nucleotide alignment under the CAT-GTR (Lartillot & Philippe, 2004) model with 10 gamma rate categories. We ran two independent runs sampling every 10th cycle and checked the convergence with Tracer 1.7.2 (Rambaut et al., 2018). The runs converged quickly and we stopped the them after collecting 3,284 and 3,113 samples, respectively. Both runs produced identical consensus trees.

**Table 2.**
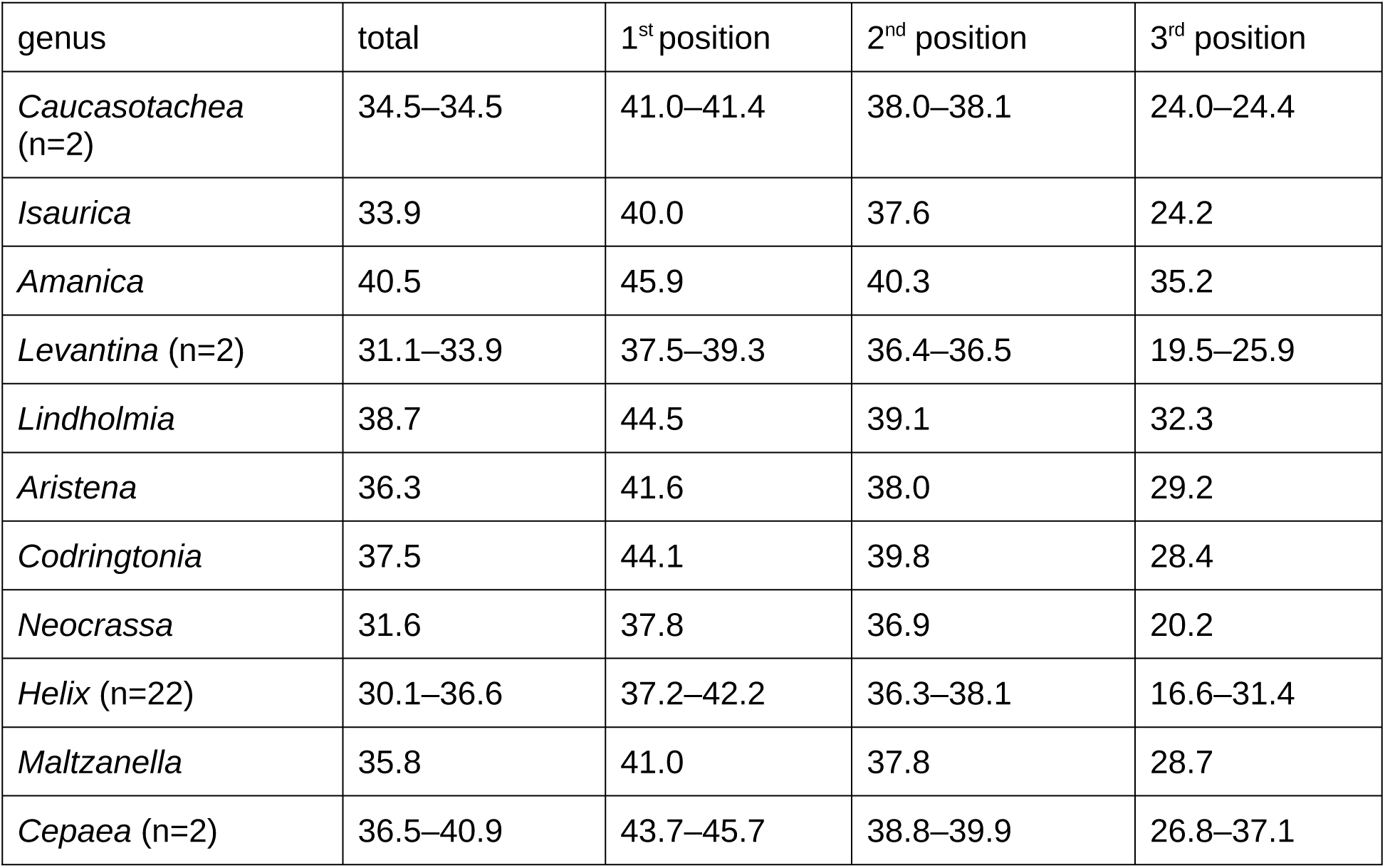
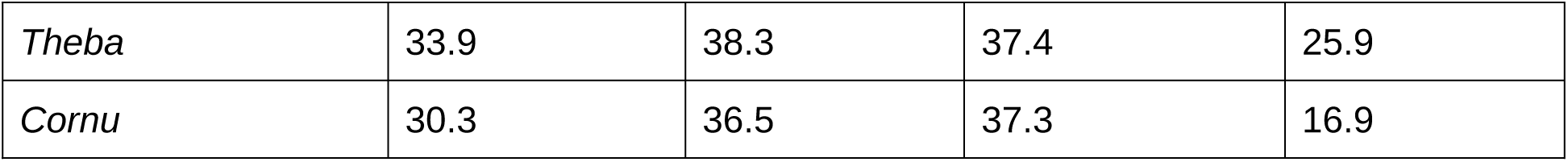
Variation in nucleotide composition among the Helicini genera and the outgroups (*Cepaea*, *Theba*, and *Cornu*). Based on a concatenated alignment of 12 protein-coding genes (*atp8* excluded).

Due to the observed heterogeneity in nucleotide composition (Table 2) and substitution rates (Table 3), we alsp explored the effects of taxon removal on the topology. We removed one genus at a time or kept only *C. aspersum* as outgroup and rerun the tree search and bootstrapping with the filtered nucleotide alignment and mixture model in IQ-TREE as above.

**Table 3.**
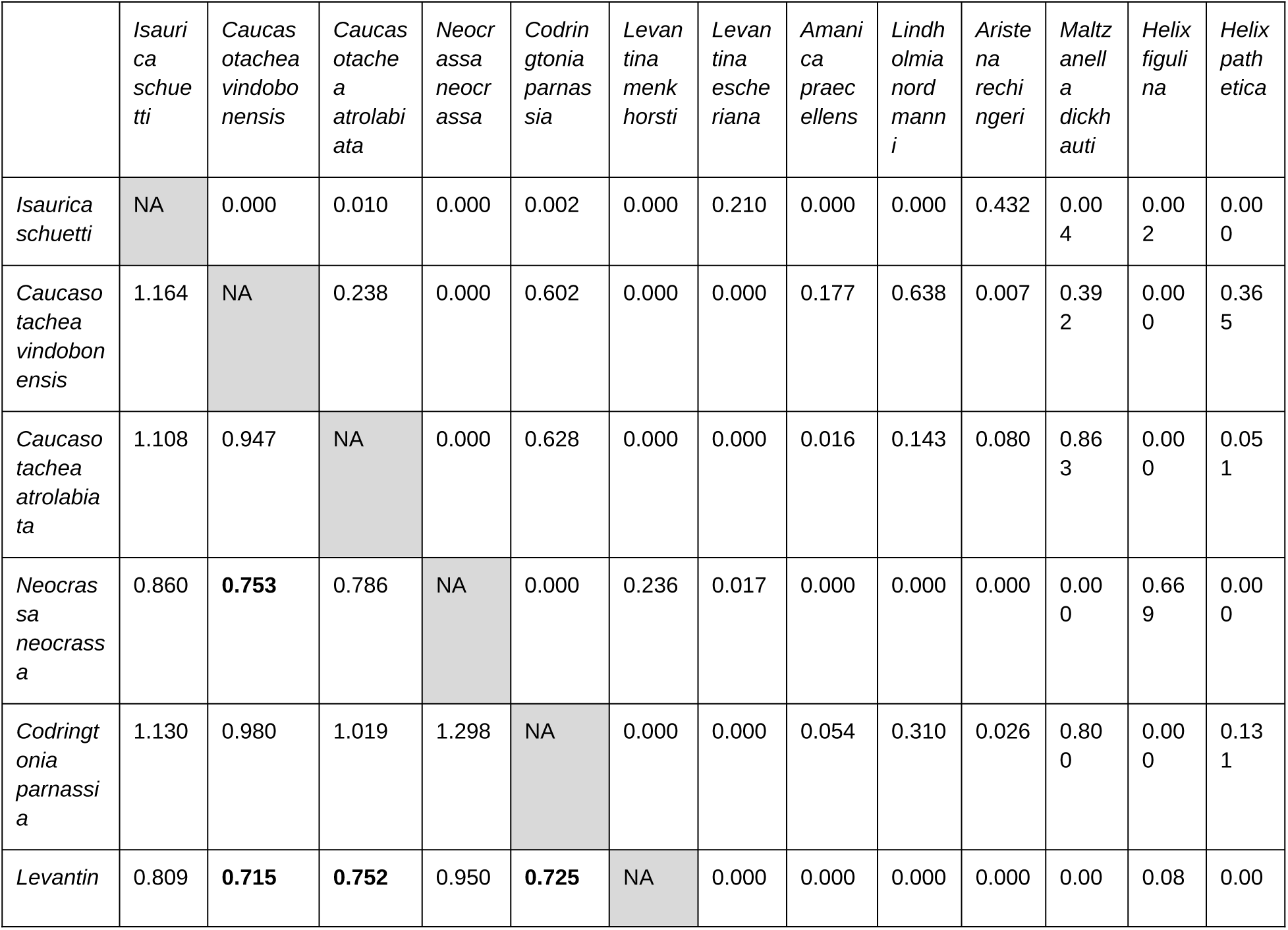
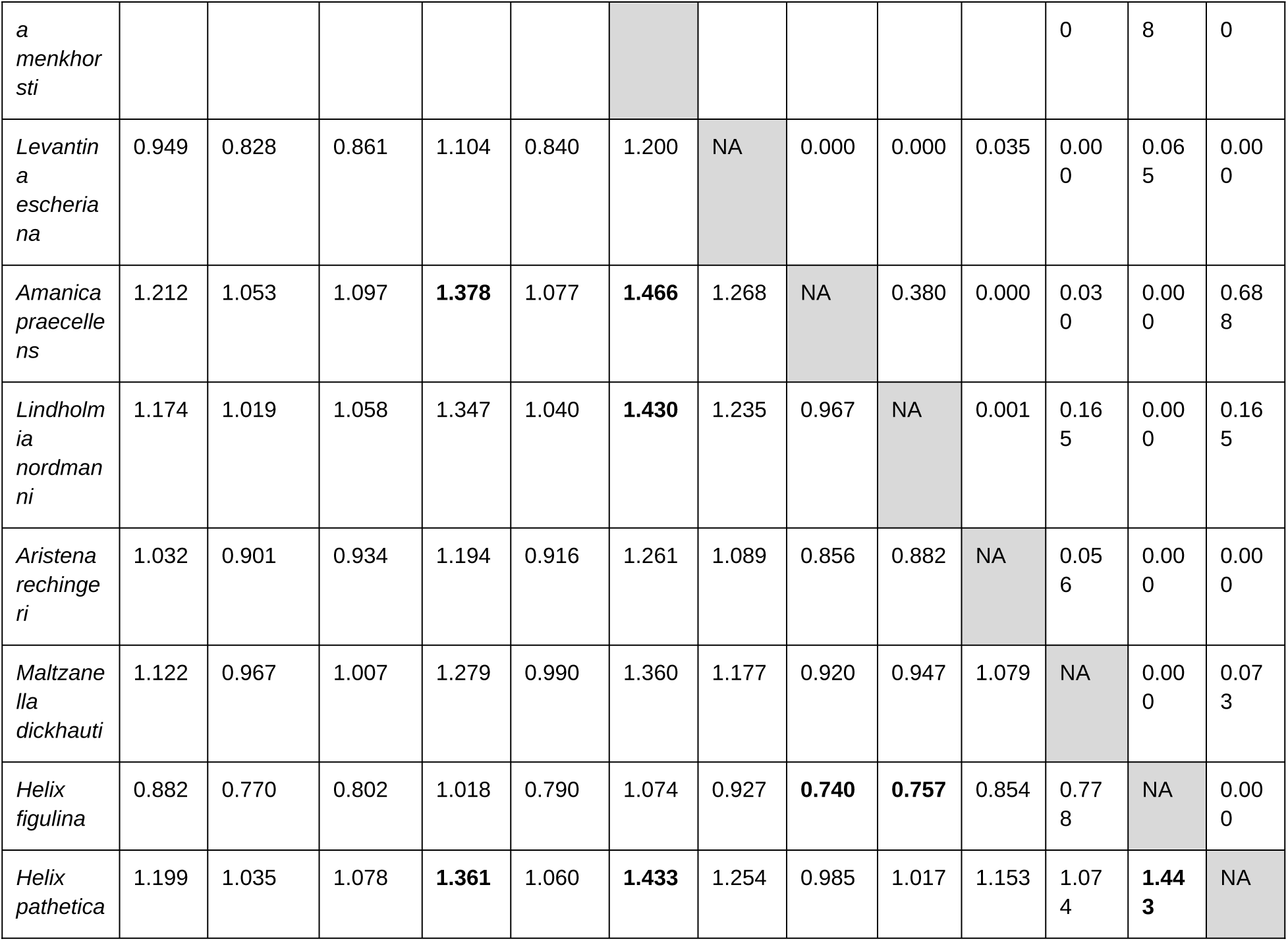
Results of Tajima’s relative rate tests performed on the unfiltered nucleotide alignment of 12 protein-coding mitochondrial genes. Above the diagonal are p-values (but note that some combinations of the tests are not independent), below the diagonal are the ratios of the numbers of substitutions unique to the lineage of the species in the row vs. of the species in the column. The most extreme differences are highlighted in bold.

### Analysis of rate variation

To explore the variation in the substitution rates independently on the topology of the phylogenetic trees, we employed the Tajima’s relative rate test (Tajima, 1993) in MEGA 11.0.13 (Tamura et al., 2021). We performed the tests on all pairs of ingroup samples except for *Helix*, where we selected the species that had either the shortest or the longest root-to-tip distance in the phylogeny. The method returns an estimate of the number of substitutions unique to each of the two tested samples, which we then used to calculate ratios that show the magnitude of rate difference.

## RESULTS

### Mitogenome assemblies

Technical statistics of the sequencing and assembly (number of raw and filtered reads, coverage, mapping rate, number of gaps in the assembly) are provided in the Supplementary Table S2. In total, 37 mitogenomes were analysed including outgroups, nine of them assembled exclusively for this study. Eight out of the nine were assembled as presumably complete mitogenomes or with only a few bp missing.

### Mitogenome characteristics

#### Gene order

Considering protein- and rRNA-coding genes, the gene order in all studied species corresponds to the order typical for Euthyneura (Varney et al., 2021) with the exception that *cox3* precedes *nd4*, which is characteristic for Helicidae (Zhao et al., 2023). The heavy strand codes for the majority of genes and is, therefore, referred to as the plus strand. All the genes coded by the light strand form a single block. We confirm that compared to *Cylindrus obtusus* (Draparnaud, 1805), the only member of Ariantinae Mörch, 1864 with a published sequence so far (Groenenberg et al., 2012), Helicinae mitogenomes differ in the positions of *trnA* and *trnP*, which are swapped. We found minor differences between the studied species in gene order concerning some tRNA genes. In *C. parnassia*, the order of *trnH* and *trnG* is reversed. *trnN* is inserted between *trnH* and *trnQ* in *I. schuetti*, while a pseudogenized *trnN* remains at its original position between *atp8* and *atp6*. In *H. pronuba*, the region involving *trnS2* is duplicated and the functional copy appears to be the one in the new position.

Genes for three of the subunits of the complex I (*nd5*, *nd1*, *nd4L*) are translated from a single mRNA. There is an overlap between *nd5* and *nd1* (10 bp in *Caucasotachea*, 13 bp in *Helix*, 14 bp in *Amanica* Nordsieck, 2017, 19 bp in *Neocrassa* Subai, 2005, *Codringtonia* Kobelt, 1898, *Aristena*, *Isaurica*, *Levantina*, *Lindholmia* Hesse, 1918, and *Maltzanella*), which is found also in other helicids (13 bp in *Cornu aspersum*, *Cepaea hortensis*, 19 bp in *Cepaea nemoralis* and *Theba pisana*, 22 bp in *Cylindrus obtusus*). There is usually also a one base overlap between the stop codon of *nd1* and the start codon of *nd4L*.

#### Start and stop codons

Alternative start codons are used in invertebrate mitochondria. We found most frequently ATG, while GTG and TTG are also frequent and CTG is used rarely (mostly in *nd6*). We found only three cases of ATA, always in *nd5*; ATT and ATC are not used.

The two genes that are followed by another protein-coding gene translated from the same mRNA (*nd5*, *nd1*) always end with a complete stop codon (TAA or TAG). In most other cases, the stop codon TAA is completed by polyadenylation of the mature transcript following the excision of tRNA(s). However, sometimes there is a full stop codon and the poly(A) tail follows after another T or NT (typical for *nd4* in *Helix*).

#### rRNAs

The exact limits of the rRNA genes are uncertain, except for the 3’ end of *rrnL* in some of the snails sequenced for their transcriptomes where the poly(A) tail was observed. The issue of the exact start and end positions dates back already to the first two land snail mitogenomes sequenced. For *Albinaria*, Hatzoglou et al., (1995) assumed that the rRNA genes occupy the whole space between the adjacent genes and Terrett et al. (1996) determined the start and end positions only provisionally based on comparison with *Drosophila*. In the reference mitogenome sequence of *H. pomatia* (NC_041247.1; Groenenberg & Duijm, 2019), the *rrnL* gene is annotated as starting right after the end of the preceding tRNA gene. That is probably correct, since the length of the stretch between the end of *trnV* and the first block within *rrnL* exhibiting clear homology between genera (the very strongly conserved TACCTTTTGCAT sequence where the 16Scs1 primer anneals) is remarkably stable despite dissimilar sequences.

The situation is different between *trnE* and *rrnS* (i.e. at the end of *rrnS*). If annotated based on alignment with the reference mitogenomes of *Homo sapiens* (NC_012920) and *Drosophila yakuba* (NC_001322), there remains a short unassigned, difficult-to-align stretch of variable length (26–60 bp) between *rrnS* and *trnE*. The exact position of the end of *rrnS* is thus uncertain.

#### Duplications, non-coding regions, control region, origins of replication

Although the stylommatophoran mitogenomes remain short and with few, short non-coding regions in the long-term evolutionary perspective, a recent study found frequent duplications in the mitogenomes and high levels of length heteroplasmy (Davison et al., 2024). Here we used short-read sequencing, which is not well suited for detection of duplicated regions, but we assembled some long duplicated or non-coding regions anyway and found additional cases where assembly failed likely because of duplications.

Long insertion was previously found between *cox1* and *trnV* in *Levantina spiriplana* (Korábek, Juřičková, et al., 2022), but this is missing from *L. menkhorsti* and largely also from *L. escheriana*. There is a long non-coding region between *trnV* and *rrnL* in *Codringtonia elisabethae* (Korábek, Juřičková, et al., 2022). This region is present also in *C. parnassia*, where *trnV* appears to be duplicated in addition. However, we are not fully confident that the assembly is complete here without long-read sequencing. *trnV* is repeated up to 12× in tandem in the *Cepaea nemoralis* mitogenome, as revealed by PacBio long read sequencing (Davison et al., 2024). The translocated *trnN* in *Isaurica* is flanked by shorter non-coding regions, as is also *trnS1* in *H. godetiana*. *trnN* in *Lindholmia nordmanni* is also preceded by ∼320 bp of unassigned sequence.

Duplications of protein-coding genes were detected in *H. pronuba* and *H. ligata*, in both cases involving genes close to the boundary between the blocks coded by the heavy and light strands, respectively. In *H. pronuba*, the region containing *trnS2, trnT* and *cox3* is duplicated, with signs of ongoing pseudogenization (frameshift deletions in the *cox3* copy). *atp8* might be duplicated in *H. ligata*, as reads from two sequence variants mapped to the assembled *atp8* gene sequence.

The locations of the control region and of the origins of replication (O_H_, O_L_) are so far uncertain. For Helicidae, it was suggested that the control region is located between *cox3* and *trnS1* (Groenenberg et al., 2012; Gaitán-Espitia et al., 2013). There is consistently a ∼180–190 bp long unassigned region in the analysed Helicini species between these two genes, located at the boundary between the blocks of genes coded by the heavy and light strand, respectively. It is the most likely location for the control region and O_H_. The light strand origin of replication (O_L_) may be located between *trnP* and *nd5*, as there is consistently a second non-coding region with a length usually exceeding 30 bp. In mammals, O_L_ is a short (∼30 bp) sequence (Sahyoun et al., 2014), so this region is probably long enough to contain O_L_. The shortest sequence between *trnP* and *nd5* was found in *C. vindobonensis* (23 bp).

#### Nucleotide composition

We examined the GC-content of the protein-coding sequences used for the phylogenetic analyses (Table 2). On the lower end, *H. figulina* and *H. pelagonesica* had a GC-content of 30.1 and 30.6%, respectively. On the high end, it reached 38.7% in *L. nordmanni* and 40.5% in *A. praecellens*. The outgroup shows a very similar range of variation. The variation differs between codon positions, with the greatest differences in the 3rd position (16.6–35.2% among Helicini, 16.6–31.4% just within *Helix*). Variation within the outgroup was not smaller even though it comprised only four species.

#### Rate variation between species

We performed Tajima’s relative rate tests on the unfiltered nucleotide alignment for all pairs of genera; for genera with two sampled species we included both of them and for *Helix* the two species displaying the shortest and longest branch, respectively. *Cornu aspersum* was used as an outgroup in all comparisons. The least evolutionary change took place along the line leading to *L. menkhorsti*, the most derived sequence is that of *A. praecellens* (Table 3). In their pairwise comparison, *A. praecellens* had ∼1.5× more unique substitutions than *L. menkhorsti*. This figure is uncorrected for multiple substitutions.

### Rate variation between PCGs

Mitochondrial protein-coding genes evolve at different rates depending on the selective pressure upon the function of the proteins they code for. As expected (see Romero et al., 2016), the most conserved gene is *cox1*; other conserved genes are *cox2* and *cox3*, a large part of *cytb*, and part of *nd5* (Figure 1).

**Figure 1.**
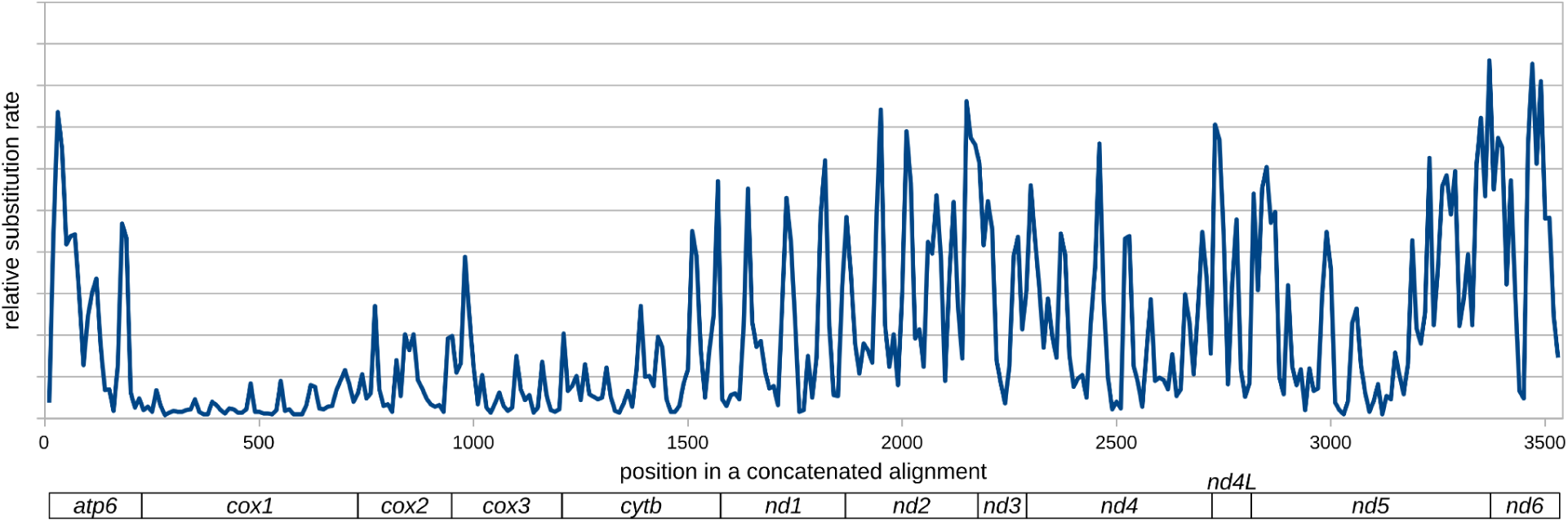
Relative substitution rates of amino acid sequences of the genes used in the phylogenetic analyses, shown as averages in 10 aa bins. Rates were estimated with MEGA 11 using a Neighbor Joining tree and mtREV+F+G6 model. The fastest-evolving gene, *atp8*, was difficult to align and was not included.

### Mitochondrial phylogeny Helicini and the position of *Aristena*

All three Maximum Likelihood (ML) analyses placed *Aristena* as the sister group to a maximally supported clade uniting *Helix* and *Maltzanella* (Figure 2A). This position was not statistically supported by the partitioned analysis (BS 68%), but received high support from an analysis of filtered nucleotide alignment with a mixture model (BS 90%). Also the Bayesian analysis, performed with the most complex substitution model, yielded strong support for this relationship (Figure 2B). Exclusion of individual ingroup genera or long-branched outgroup taxa did not change this grouping, with bootstrap support from the mixture model in IQ-TREE varying between 83% and 91% with the exception when *Maltzanella* was excluded, which led to a diminished support (BS 61%).

**Figure 2.**
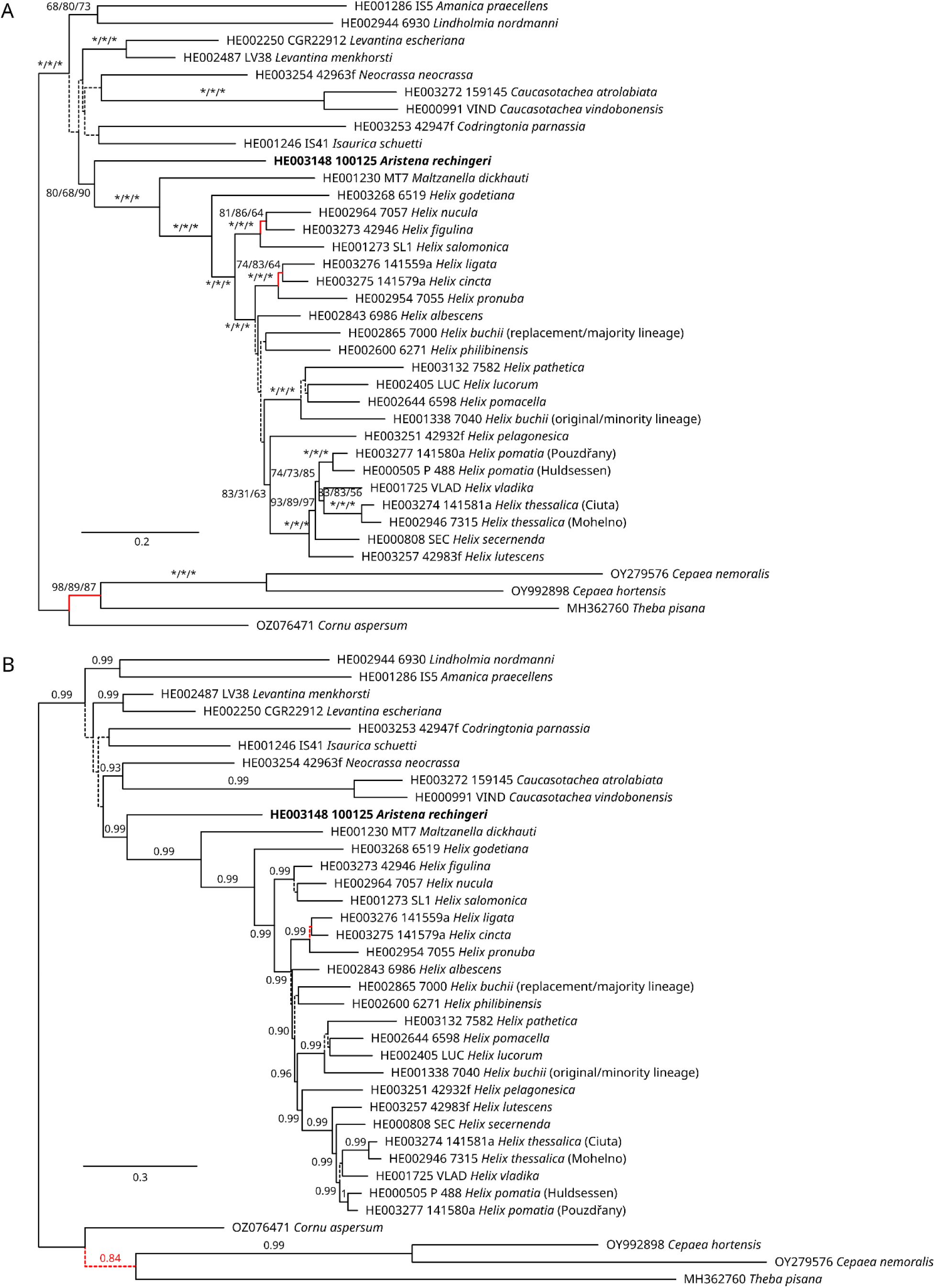
Mitochondrial phylogeny of the tribe Helicini showing the position of *Aristena rechingeri*. A, Maximum Likelihood tree from an analysis with a mixture model of on alignment of nucleotide sequences of 12 protein-coding genes (excluding *atp8*) filtered by ALISCORE and with third codon positions removed. Branches with bootstrap support from that analysis <50% are shown as dashed lines. The support values derive from (in this order) a partitioned analysis of the unfiltered nucleotide data, partitioned analysis of a translated amino acid alignment and from the analysis of filtered nucleotide data with a mixture model. Asterisks denote 100% bootstrap support. B, phylogeny inferred with PhyloBayes using the CAT-GTR+Γ10 model on the filtered nucleotide alignment as in A. Dashed branches indicate posterior probabilities <0.90. Presumably incorrectly resolved reference branches, which are most likely an artefact of long-branch attraction/repulsion (see Methods) are shown in red.

Apart from the clade of *Helix*, *Maltzanella*, and *Aristena*, the only other supported grouping among genera is a clade uniting *Lindholmia* and *Amanica*, but with a weaker support in the ML analyses. The support varied a bit with the exclusion of individual genera from the mixture model ML analysis (BS 72–86%, 60% if *Helix* excluded). The PhyloBayes analysis strongly supported this grouping.

There were three points in the tree where we had prior expectations on what the correct topology should be. In two clades of *Helix*, the expectations stemmed from a multilocus nuclear phylogeny and mitochondrial phylogenies inferred with different substitution models (Korábek & Hausdorf, in prep.). The expected topology in the outgroup followed previous analyses of short mitochondrial sequences from a broader set of related species and of nuclear ITS2 sequences (Koene & Schulenburg, 2005; Razkin et al., 2015; Korábek, Juřičková, et al., 2022; Neiber et al., 2022). In all cases the ML recovered here a topology considered incorrect. The analysis of filtered alignment with mixture models resulted in lower support values than analyses of unfiltered nucleotide or amino-acid data with partitioned models, but still the incorrect grouping of *Theba* and *Cepaea* received significant support (BS 87%). The Bayesian inference (BI) with PhyloBayes was a bit more successful, recovering the correct placement for *H. figulina* (without significant support) and low support for the incorrect groupings of *H. cincta* and *H. ligata* (PP 0.52) and of *Cepaea* and *Theba* (PP 0.84). PhyloBayes also produced a tree with greater differences in root-to-tip distances between species than ML.

#### Isaurica callirhoe

The only finding of *I. callirhoe* after its description was reported by Ceylan et al. (2008). Here, we sequenced a preserved individual from that study. The results of phylogenetic analysis of mitochondrial data put *I. callirhoe* clearly within *I. lycia* as currently accepted (Figure 3), close to samples from Megisti Isl. (Kastellorizo Isl.). There are no real conchological or anatomical differences between *I. callirhoe* and *I. lycia* from Megisti (Supplementary Appendix S2).

**Figure 3.**
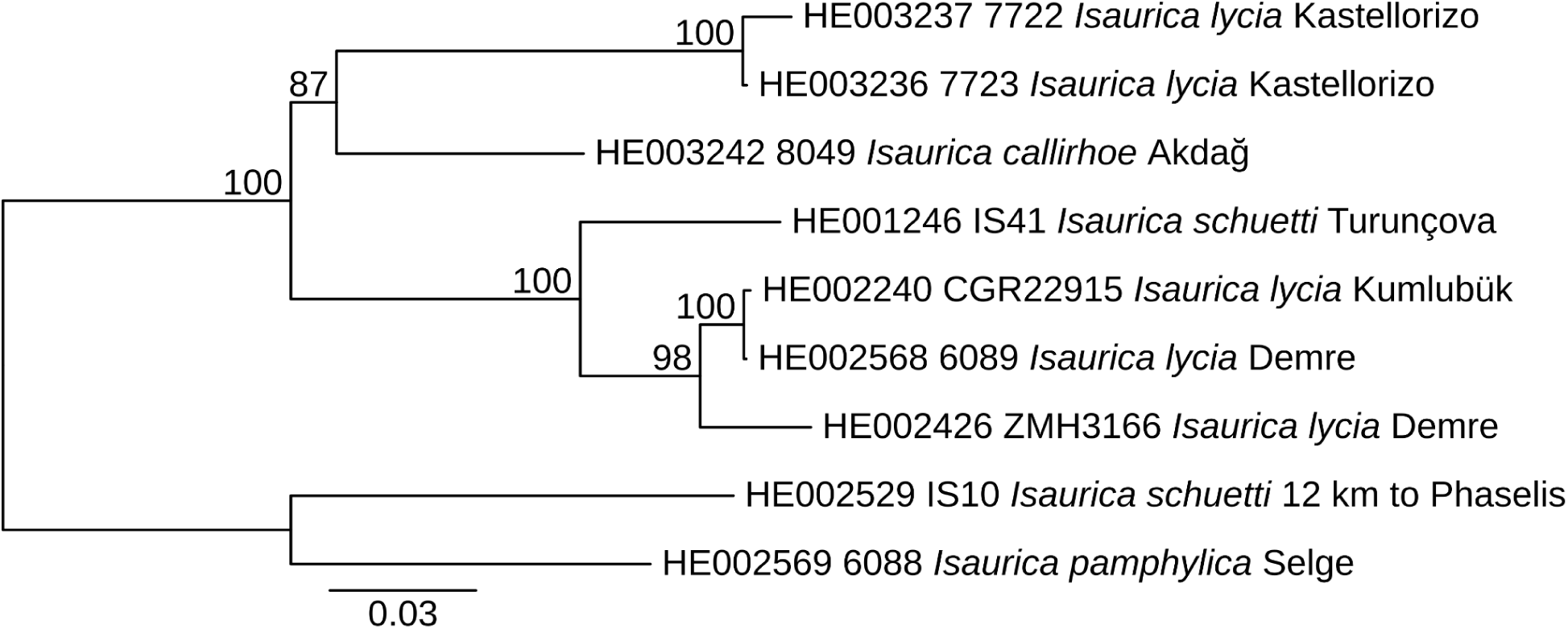
Phylogenetic relationships within *Isaurica*. Maximum likelihood tree based on concatenated mitochondrial nucleotide sequences of partial *cox1*, *rrnL*, and *rrnS* genes. Support values are bootstraps from 500 pseudoreplicates. No outgroup was included in this analysis as there is no one closely related, but the root position is as implied by the tree layout (Korábek, Juřičková, et al., 2022).

The two analysed samples of *I. schuetti* were each placed elsewhere in the tree. The one that we used for mitogenome assembly was found nested within *I. lycia*. We suggest that this is due to a mitochondrial introgression from *I. lycia*, as the sample was collected near the boundary between the distribution range of the two species.

## DISCUSSION

### The closest relatives of *Aristena* and the phylogeny of Helicini

In agreement with the results of Psonis et al. (2022), *Aristena* was found here to be a sister lineage to *Helix* + *Maltzanella*. In turn, we found no support for a sister relationship with the conchologically much more similar and geographically likewise close genus *Isaurica*. We also verified that the poorly known taxon *I. callirhoe* belongs to *Isaurica*; in fact, we propose to classify it as a junior synonym of *I. lycia* (see below). The sister relationship of *Aristena* to *Helix* + *Maltzanella* gained support with the use of mixture substitution models coupled with third codon position removal, and it is compatible with nuclear ITS2 phylogeny (Korábek & Hausdorf, 2023). Furthermore, it makes sense geographically. *Maltzanella* diversified in the western Anatolia, while *H*. *godetiana*, a species sister to all other *Helix* species, lives on islands in the Aegean, and the crown group of *Helix* likely originated from the Aegean or western Anatolia (Korábek & Hausdorf, in prep.). *Aristena*, living on an island in the southeastern Aegean, is thus a geographically likely candidate for a sister group of *Helix + Maltzanella*.

*Aristena* is apparently a relic taxon, containing only one species that occurs on a single island and which is very rare. As follows from our results, it uniquely documents the geographic origins of *Helix*, one of the most diverse and most broadly distributed helicid genera. Another such relic species from the Aegean, *H. godetiana*, is in a long-term decline and appears to be under a threat of extinction by the increasingly dry climate due to the timing of its reproduction period (Maroulis et al., 2025). Studies of physiology of aestivation suggest that *H. figulina* and *Codringtonia helenae* (from Lesvos and Peloponnese, respectively) might struggle if the aestivation period becomes longer due to climate change (Giokas et al., 2007; Kotsakiozi et al., 2012). Besides mapping the surviving populations, it is now important to investigate the biology *A. rechingeri* to see if it is facing similar threats.

Unfortunately for the resolution of the mitochondrial phylogeny, there are short branches between the Helicini genera followed by long branches to the tips, suggesting a fast diversification. This was apparent already from earlier analyses with partial sequences of a few genes (Korábek, Juřičková, et al., 2022; Neiber et al., 2022). Not surprisingly, the analyses of both the nucleotide and amino-acid sequences resulted in an unresolved base of the tree with very low support values. Besides the grouping of *Maltzanella* and *Aristena* with *Helix*, the only other supported relationship between genera was a clade comprising *Amanica* and *Lindholmia*. Here the support was lower than in the case of *Aristena*, but the clade was also already suggested by earlier analyses (Korábek, Juřičková, et al., 2022; Neiber et al., 2022). Both belong to the three eastern-most Helicini genera (the third and easternmost one being *Levantina*; Korábek, Glaubrecht, et al., 2022), and although their ranges are widely geographically separated, the gap may be just another case of separation of related land snail taxa caused by the uplift and aridification of the Anatolian Plateau (Neiber, Walther, et al., 2021; Lehmann et al., 2024). The two genera have the highest GC content of all Helicini, so their grouping may theoretically be an artefact caused by that; however, the higher GC content may as well be a synapomorphy of the group.

The incorrect topology of the three reference subclades inferred in our analyses (red branches in Figure 2) show that the analyses struggled to find the true tree. Nevertheless, the support for the presumably incorrect branches decreased from the unfiltered nucleotide data and site-homogeneous partitioned model to the filtered data and site-heterogeneous mixture model, being the lowest with the most complex CAT-GTR model, while the support increased for the two above groupings of interest, lending the results some confidence. Analyses of filtered nucleotide alignments with mixture models also yielded the most credible trees in the case of *Helix* (Korábek & Hausdorf, in prep.).

### The status of Isaurica callirhoe

*Isaurica callirhoe* was described in 1894 from a locality imprecisely described as “Lycia, in montibus Ak-Dagh nec non prope Macri” (Rolle, 1894). As explained in Rolle & Kobelt (1896), it was collected while travelling across the mountains from Fethiye (formerly Makri) to Elmalı; the label of the original series specifies the locality as the northern margin of Akdağ. As the map in Subai (1994) suggests, the area could be part of the range of *Isaurica lycia*, from which *I. callirhoe* should differ in having a flatter shell with granulated upper surface (due to fine spiral grooves), narrower and paler bands, and a lower aperture (Rolle, 1894; Subai, 1994). There is no close relationship between *I. callirhoe* and *A. rechingeri*, and we can also reject Nordsieck’s (2017) speculations that *I. callirhoe* is in fact a species of *Levantina*. The mitochondrial phylogeny of *Isaurica* (Figure 3) places *I. callirhoe* within the intraspecific diversity of *I. lycia* as delimited by Subai (1994). *Isaurica callirhoe* is not as conchologically different from *I. lycia* as claimed by Subai (1994), because the shape and colouration of the latter are variable and not all populations have such high shells as the individual depicted by Subai. Especially the westernmost populations, like those from Megisti Isl., have much flatter shells and a slightly different colouration, matching in appearance *I. callirhoe*. There may also be some spiral grooves on the surface (see also the figures in Pfeiffer & Wächtler (1939).

In light of the above, we consider *I. callirhoe* a synonym of *I. lycia*.

### Heterogeneous evolution of land snail mitogenomes and phylogenetics

Besides short internal branches, there are other reasons why reconstructing the phylogeny of mitochondria in Helicini is difficult. Our study documented variation in parameters of the mitogenome evolution that are likely to affect phylogenetic analyses.

There is an obvious variation in the substitution rates as demonstrated by the Tajima’s test, which is independent on details of the topology (but ignores multiple substitutions per site) as well as by the substantial variation in root-to-tip distances (as seen especially in the PhyloBayes analysis; Figure 2B). Another issue, besides the rate heterogeneity, is heterogeneity in the nucleotide composition. The difference between the lowest and the highest GC content in the protein-coding sequences among species exceeds 10 percentage points in both the ingroup and the outgroup. This may also be relevant for the phylogenetic analyses. The situation resembles that documented previously well with insect mitogenomes (e.g., Timmermans et al., 2016; Liu et al., 2018, and references therein), where the heterogeneity also led to incorrect groupings. As was also the case here, the analyses with site-heterogeneous mixture models produced more defensible trees for those insect taxa. However, in our case the phylogenetic depths considered are much shallower than in these examples.

In the ingroup, the position of *H. pronuba* (long branch, high GC content) relative to *H. cincta* and *H. ligata* (short branches, low GC content; Figure 2; Korábek & Hausdorf, in prep.) was affected by rate difference, difference in nucleotide composition, or both. However, the problems with long-branched taxa, some of them with unusual GC content, can be best demonstrated here in the outgroup. Three out of the four available species, including *Cepaea nemoralis*, have very long branches, i.e. accelerated substitution rates. The fourth one, *Cornu aspersum*, has a markedly shorter branch. Furthermore, the GC content of *C. nemoralis* is the highest in the dataset, while that of *C. aspersum* is among the lowest and *T. pisana* has a higher GC content than *C. aspersum*. In all analyses the long-branched *Theba*, which is more closely related to *Cornu*, grouped with the likewise long-branched *Cepaea*. With the sparse sampling here, this incorrect grouping appeared in all analyses and only the most complex one with PhyloBayes did not find statistical support for it. The phylogenetic inference of the relationships within the outgroup could be improved with additional sampling, because analyses of much shorter alignments with site-homogeneous partitioned model, but with much more complete sampling of genera, were able to recover relationships between *Cornu*, *Theba* and *Cepaea* compatible with the nuclear phylogeny (e.g. Korábek, Juřičková, et al., 2022). Unfortunately, additional sampling is not an option for improving the resolution of the relationships of the monotypic or species poor genera studied here. Furthermore, the negative effects of the compositional heterogeneity are expected to be the worst with short branches separating long branched taxa, as is the case with Helicini (Jermiin et al., 2004).

The observed variation between relatively closely related species shows that the evolution of mitochondrial genome sequences in land snails is highly uneven in pace and direction. A recent analysis of *C. nemoralis* found widespread heteroplasmy and length variation (Davison et al., 2024), suggesting unusually fast evolution of its mitogenome, a hypothesis supported here. However, despite sometimes very fast evolution, the land snail mitogenomes remain similar in length, gene order (Zhao et al., 2023) and retain function over long evolutionary timeframes. We observed some duplications and subsequent pseudogenizations, resulting nevertheless only in minor changes to gene order affecting the position of tRNA genes. Species and possibly also genera with fast and slow substitutions are sometimes closely related, showing that the speed of evolution changes quickly back and forth. The GC content varies, but within the same limits in the ingroup as in the outgroup. We suggest that while these variations may cancel out in the long term, they may cause mitochondrial sequences to be difficult data for phylogenetics exactly where these are used the most, at the species and genus levels.

### Annotation of land snail mitogenomes

Our representative sampling of mitogenomes across Helicini, a subclade of the land snail family Helicidae, allowed us to improve the annotation of Helicid mitogenomes. This is an issue that has been mostly neglected, so despite the number of available mitogenome sequences of land snails, several unaddressed issues remain.

First, to correctly annotate land snail mitogenomes, it is necessary to consider all possible alternative start codons. Ghiselli et al. (2021) listed ATG, TTG, GTG, ATA, ATT, and ATC as possible start codons in molluscs and CTG was also reported (Wu et al., 2010). By aligning and comparing sequences of a number of related species, we found genes starting at congruent positions with ATG, GTG, TTG, CTG and ATA, although the last two are used rarely. In *H. pomatia* from Huldsessen we identified only ATG, TTG and GTG as start codons, but automatic annotation with MITOS2 correctly identified only six start codons (all ATG). For comparison, MitoZ 3.6 (Meng et al., 2019) found eleven (although for the genes on the minus strand the positions were off by a single bp).

Second, the stop codons appear to be usually completed only with the polyadenylation of the transcripts. This is a phenomenon long known from mammals (Anderson et al., 1981), but insufficiently appreciated for land snails. Complete stop codons in the species analysed here are only consistently present in the two genes that are followed by another protein coding gene translated from the same mRNA. Protein-coding genes followed by tRNA typically end with an incomplete stop codon completed by polyadenylation of the mature transcript. This contrasts with the conclusions of Groenenberg & Duijm (2019), who claimed a full stop codon for each of the 13 protein-coding genes of the *H. pomatia* mitogenome. It is, thus, necessary to correct the outputs of automatic annotation tools, which search for stop codons in the mitogenome sequences and take into account existing annotations (Donath et al., 2019; Meng et al., 2019). In our annotation, 10 of the protein coding genes of *H. pomatia* end with an incomplete stop codon, while it is only 5 in the RefSeq NC_041247.1. MITOS2 correctly identified one incomplete stop codon in the sequence of *H. pomatia* from Huldsessen (*cox3*), while MitoZ 3.6 (Meng et al., 2019) identified two (*atp6*, *cox3*).

Third, the exact limits of genes coding for tRNAs and rRNAs cannot be located based on alignment only. The annotation of tRNA ends by MITOS 2 (Donath et al., 2019) is sometimes inconsistent between species. Finding the exact RNA gene limits would require targeted analyses beyond the scope of this work.

Fourth, the location of the control region and the origins of replication remains uncertain. Several studies speculated on the possible location of the control region in land snails (Hatzoglou et al., 1995; Deng et al., 2016; He et al., 2016; Inäbnit et al., 2024), but none provided conclusive evidence. We conclude that the location suggested by Gaitán-Espitia et al. (2013) is likely correct.

### Re-evaluation of *Levantina menkhorsti* (Gümüş & Neubert, 2012)

As a byproduct of our sampling for the analysis of mitogenomes, we revise here the taxonomic status of *Levantina menkhorsti* (Gümüş & Neubert, 2012). We sequenced two samples of *Levantina*, separated by the deepest split within the genus. One of them, namely LV38, had not been identified to species level previously (Korábek, Glaubrecht, et al., 2022). We identified it now as *L. menkhorsti*.

Described as a subspecies of *Levantina thospitis* (Schütt & Subai, 1996), *L. menkhorsti* is the latest contribution to the list of *Levantina* species-group taxa. It was described from shell material originating from two sites in the Bitlis Province of Turkey (south of Bitlis and in the uppermost valley of Kezer Çayı south of Tatvan; Gümüş & Neubert, 2012). This is only 21 and 7 km, respectively, from Küçüksu, where the sample LV38, analysed here, was collected. The shells from Küçüksu shared one of the defining characteristics of *L. menkhorsti*, the greenish-yellowish periostracum, and their shell shapes match perfectly.

We consider *L. menkhorsti* to be a species distinct from *L. thospitis*. One reason is the divergence in mitochondrial sequence, as the mtDNA of *L. menkhorsti* is basal to mtDNA of all other *Levantina* species (Korábek, Glaubrecht, et al., 2022). Another reason is that the conchological differences between the two are, in the context of the genus, relatively strong. Finally, the collection of H. Schütt in SMF contains samples from an area in which both *L. menkhorsti* and *L. thospitis* come in contact and where they appear as clearly distinct morphologically with the exception of a few intermediates. The contact zone is located in the Batman Province of Turkey near Kozluk (Hezzo); samples were collected by H. Schütt on 02.– 03.05.1999 and 26.05.2001 at two mutually close sites in the valley of Yanarsu (Garzan) Çayı and the material is deposited in the Senckenberg Natural History Museum, Frankfurt/Main, Germany (coll. Schütt 1571 and 1577a).

*Levantina thospitis* differs from *L. menkhorsti* in having a more expanded last whorl with a spacious aperture that has a strongly broadened, reflected lip. *Levantina thospitis* also lacks the greenish-yellowish periostracum of *L. menkhorsti*. The colouration differs in that the bands have a blurred appearance in *L. thospitis*. Furthermore, *L. thospitis* has an always closed and thickly calloused umbilicus (often slit-like open in *L. menkhorsti*) and smoother shell surface with weaker ribs.

## CONCLUSIONS

We used mitogenome sequences to resolve the phylogenetic position of an extremely rare single-island endemic, *Aristena rechingeri*. We found no support for a relationship to the geographically closest flat-shelled genera *Codringtonia* or *Isaurica*, but we were able to confirm that the flat-shelled *Aristena* is the sister group of the globular-shelled genera *Helix* and *Maltzanella*. However, the mitogenome sequences turned out to be difficult data for phylogenetic analysis. Malacologists focusing on the systematics of land snails have for years been largely ignorant of the patterns of mitogenome evolution despite relying heavily on mitochondrial sequences in their work (see Böckers et al., 2016 for an exception). Several aspects of the mitogenome evolution in land snails thus remain understudied or disregarded. Substitution rates and nucleotide compositions vary substantially between related species, and reconstructing the mitochondrial phylogeny may be very difficult even with complete mitogenomes. Heterogeneity in the substitution process among branches remains the most difficult problem for phylogenetic inference (e.g. Kapli et al., 2023) and, along with the study of Korábek & Hausdorf (in prep.), we conclude here that the usual approach, i.e. to perform a partitioned analysis with a site-homogeneous model, may lead to false conclusions even at the shallow phylogenetic depths examined here. As regards the mitogenome structure, duplications in the mitogenomes are not rare, as recently also shown by Davison et al. (2024) for *Cepaea nemoralis*, but they may be rather quickly eliminated. We also found that annotations need to take into account that the majority of protein coding genes end with an incomplete stop codon and that there are several alternative start codons. It is therefore necessary to use alignments to look for start and stop positions that would be consistent between species rather than relying on existing annotations.

## SUPPLEMENTARY DATA

**Appendix S1.** Detailed description of the methods used to sequence and assemble the mitogenomes analysed here (other than *Aristena rechingeri*).

**Appendix S2.** Genital system anatomy of *Isaurica callirhoe* and *I. lycia*.

**Table S1.** List of the samples analysed along with locality and voucher information and accession numbers of the sequences deposited in the NCBI’s nucleotide (assembled mitogenomic sequences) and SRA (raw reads) databases.

**Table S2.** Technical statistics concerning the sequencing and assembly results of the mitogenomes analysed here. See text for methods of quality filtering, which differed between batches. Coverage was calculated from filtered reads mapped onto the final assembly with Bowtie2 (WGS: --end-to-end -N 1 -L 20 --no-discordant; RNA-seq: --local -N 1 -L 20 --no-discordant). Note that the extremely high coverage maxima for the RNA-seq data are due to massively overrepresented *rrnL*. The lower limits of the coverage ranges marked with an asterisk are lowered by underestimation at the ends where the sequence was linearised (although 235 bp sequences were copied to each end).

## AUTHOR CONTRIBUTIONS

Conceptualization, data curation, formal analysis, funding acquisition, resources, visualization, writing—original draft preparation O.K.; methodology, O.K., N.P.; investigation, writing—review & editing, O.K., K.V., A.Ö., M.Z.Y., N.P. All authors have read and agreed to the published version of the manuscript.

## ACKNOWLEDGEMENTS

Computational resources were provided by the e-INFRA CZ project (ID:90254), supported by the Ministry of Education, Youth and Sports of the Czech Republic.

We thank Lucie Juřičková, Kateřina Kubíková, Petr Dolejš, and Radovan Coufal (Praha, Brno) for help with sampling, Ira Richling (Stuttgart) for samples of *Isaurica*, and Sigrid Hof (Frankfurt) for access to the collections of the Senckenberg Natural History Museum. Mehmet Z. Yıldırım helped to collect the sample for *Isaurica callirhoe*. Petr Svoboda (Praha) and Bernhard Hausdorf (Hamburg) provided resources for sequencing.

## DATA AVAILABILITY

Assembled mitochondrial sequences are available from the NCBI nucleotide database; accession numbers are provided in Table 1. Raw Illumina reads are available from the NCBI Short Read Archive database; accession numbers are provided in Table 1.

