## Appendix 1 for "Finding the relatives of a critically endangered island endemic reveals a heterogeneous evolution of land snail mitogenomes (Gastropoda: Helicidae)"

SUPPLEMENTARY APPENDIX S1

### Sequencing and assembly of mitogenomes of other studied species

*Helix pomatia* (DE)

The mitogenome was amplified by PCR in several overlapping fragments, which were then sequenced using stepwise Sanger sequencing. The methods are described in greater detail in (Korábek et al., 2019). The sequence is very similar to the complete mitogenomes published by (Groenenberg & Duijm, 2019) and was used as a reference for the other species.

*Helix lucorum*, *H. secernenda*, *H. vladika* & *Caucasotachea vindobonensis*

Mitogenomic sequences were assembled from 75 bp single-end Illumina RNA sequencing performed in 2015 (Korábek et al., 2019). RNA was extracted from fresh posterior foot tissue. Geneious R7.1 (Kearse et al., 2012) was used for adapter and quality trimming.

*De novo* assembly was performed. We used FastQC 0.11.9 (Andrews, 2010) to check the overall quality of the reads. Then, we run rCorrector 1.0.4 (Song & Florea, 2015) to correct likely erroneous *k*-mers and removed reads flagged with "unfixable_error" flag. Remaining adapter sequences and low quality bases were removed with TrimGalore 0.6.6 (Krueger, 2020) using Cutadapt 3.5 (Martin, 2011) and the following options: `--length 36 -q 20 --stringency 1 -e 0.1`. *De novo* assembly was then performed with Trinity 2.11.0 (Grabherr et al., 2011). Mitochondrial contigs were extracted from the assembly with blastn (BLAST 2.12.0+) (Camacho et al., 2009) using as query fragments of protein coding genes assembled earlier by mapping reads against *Cornu aspersum* mitogenome in Geneious (Korábek et al., 2019). The retrieved contigs were aligned to the *H. pomatia* mitogenome, visually inspected and manually joined where overlapping. For all samples, additional shorter contigs with alternative sequence were assembled by Trinity for part of the *rrnL* (16S rRNA) gene, which was hugely overrepresented in all the polyA selected RNAseq libraries used here. These contigs were not used for the final assembly except for the 5' end in *H. secernenda*, which was not present in the major contig in *H. secernenda*. In addition, variant contigs differing in the length and sequence were assembled for the region between *cox3* and *nd4* genes in *H. lucorum*, probably due to a repeat in this region, which was also excluded.

To explore the quality of the assembly, we used Bowtie2 2.4.2 (Langmead & Salzberg, 2012) (`--sensitive-local -L 10 -N 1`) to map reads to the assembled and joined contigs, visualised the mapped reads with UGENE 40.0 (Okonechnikov et al., 2012) and visually searched for assembly problems (partially masked reads, incorporation of reads with multiple mismatches, lack of overlap between reads, etc.), especially at gene boundaries and in the tRNA genes that display low coverage. Consensus sequences of the mapped reads were also exported using SAMtools 0.1.19 (Li et al., 2009) as implemented in UGENE and carefully compared to the assemblies. Only very minor errors were uncovered and corrected manually after inspection of the mapped reads.

By searching for overhanging reads and repeated mapping, we were able to iteratively close two gaps between protein-coding genes within the *H. lucorum* sequence. Mapping showed huge per-base coverage in the *rrnL* gene (exceeding seven million in case of *H. vladika*), but, as expected, low coverage for some of the positions within tRNAs or non-coding regions (as low as 2×).
We attempted to amplify and sequence the region between *cox3* and *nd4* genes in *H. lucorum* using a pair of primers that we designed to anneal to these genes (luc_POR_f: TGTACCAAACCGCAACCACA, luc_POR_r: TGTACTCGACAGTGCGAAGG). However, the PCR produced a broad smear around a weak band and, hence, no sequenceable product.

*Helix thessalica* (CZ), *H. lutescens*, *H. pelagonesica*, *Neocrassa neocrassa,* and *Codringtonia parnassia*

Complete or nearly complete mitogenomes were assembled from transcriptome Illumina sequencing reads. RNA was extracted from posterior foot tissues preserved in RNAlater. RNA sequencing libraries were prepared with the NEBNext Ultra II RNA Library Prep Kit for Illumina (New England Biolabs). mRNAs were enriched with Oligo(dT) beads, fragmented, transcribed to cDNA, and enriched by limited-cycle PCR. The libraries were sequenced on the NovaSeq 6000 platform (paired-end, 150 bp).
For *H. thessalica*, the sequencing results were delivered already trimmed of adapters and low-quality bases, whereas the other four datasets were delivered untrimmed (raw reads). We checked the trimming of *H. thessalica* data by running fastp 0.23.1 (Chen et al., 2018) under default settings except for `--detect_adapter_for_pe`. Then we run rCorrector on all datasets to correct likely erroneous k-mers and removed reads flagged by rCorrector with “unfixable_error” flag. Then we performed quality and adapter filtering with fastp under the following settings: `--detect_adapter_for_pe --cut_front --cut_tail --cut_window_size 4 --cut_mean_quality 25 --qualified_quality_phred 28 --unqualified_percent_limit 40 --n_base_limit 5 --length_required 30` (fastp also removes reads lacking mates after the previous filtering).
*De novo* assembly was then performed with Trinity. Mitochondrial contigs were found using BLAST and sequences of mitochondrial genes of *H. pomatia* as a query. They were aligned to the *H. pomatia* mitogenome and, where possible, collapsed into a single sequence covering the mitogenome. The assemblies were verified by comparison with the consensus of reads mapped on the assembled sequence with Bowtie2 as described above. The mitogenome of *N. neocrassa* was not assembled as complete due to one misassembled non-coding region.

*Levantina menkhorsti*, *Isaurica schuetti*, and *Amanica praecellens*

The sample of *Isaurica schuetti* was a dried body of a snail that was found dead in the shell; *Levantina menkhorsti* and *Amanica praecellens* were sampled from ethanol-preserved (presumably ~70%) material collected in 2002 and 1994, respectively. The DNA was extracted with a column-based kit (Tissue Genomic DNA Mini Kit, Geneaid).
All three samples were initially sequenced on a single Illumina MiSeq run (paired-end, 75 bp), but that was sufficient only for the *Levantina* sample, so the libraries from *A. praecellens* and *I. schuetti* were resequenced, sharing the second run in the ratio 1:2. The raw reads were trimmed and quality filtered with Cutadapt 4.6 using TrimGalore (`--length 30 -q 20 --stringency 1 -e 0.1`) and deduplicated with fastp; reads from the two sequencing runs were then combined.

MITObim was used for the assembly with `--kbait 21`. In the first instance, we provided fragments produced by Sanger sequencing (*rrnL*, *rrnS*, *trnV*, *cox1*, *cox2*, *cytb*; the latter two were not available for *Isaurica*) separated with Ns in place of the missing parts of the mitogenomes as the reference. This was sufficient for the assembly of the L. *menkhorsti* mitogenome, but large portions of the mitogenomes of both *I. schuetti* and *A. praecellens* were missing, necessitating the addition of more reference sequences for the baiting step. We used BWA (bwa-mem, `-k 5 -r 1 -A 1 -B 1`) to map reads on the mitogenomes of *H. pomatia* and used the consensus of the mapped reads as reference in another MITObim run.

The above approach, and in case of *I. schuetti* repeating this procedure with the *N. neocrassa* mitogenome as reference, helped to largely complete the assembly of *A. praecellens* and eliminate most gaps in the case of *I. schuetti*. However, in *I. schuetti* two large gaps remained: one containing most of *atp8* and the other within *nd2*. In *A. praecellens*, a small gap in *nd5* remained. In all three cases, there is low average nucleotide identity among genera. We tried mapping the reads to *atp8* and *nd2* sequences of the previously assembled samples, which resulted in producing a single hit for *atp8* when the sequence of *H. pelagonesica* was used as a query. Adding this read to the assembly and then using the latter as reference for MITObim with `--kbait 15` produced the complete *atp8* gene. The final gaps were closed by mapping reads using bwa-mem with the same settings as mentioned above on the incomplete assemblies.

Examination of reads mapped on the resulting assemblies revealed three misassembled places in the *Isaurica* mitogenome. Two were easy to correct by manually re-aligning the reads, but the third required a combination of iterative extension by multiple rounds of bwa-mem mapping followed by extension using MITObim.

*Helix pomacella* and *H. pathetica*

Sequencing libraries were prepared from DNA extracted from ethanol-preserved samples with a isopropanol precipitation-based protocol (Scheel & Hausdorf, 2012; Sokolov, 2000) and sequenced on the Illumina NovaSeq X platform (paired-end, 150 bp). The reads were trimmed, filtered and deduplicated with TrimGalore (`--length 140 -q 20 --stringency 1 -e 0.1`) and fastp (`--detect_adapter_for_pe --qualified_quality_phred 15 --unqualified_percent_limit 40 --n_base_limit 5 --length_required 15 --dedup --trim_poly_g`). The filtered reads were used for assembly with MITObim (`--kbait 31`), for which the *cox1* barcode sequences of both individuals were provided as reference for the initial mapping. Both samples produced complete mitogenome sequences after 29 rounds of iterative mapping, which were manually circularised.

*Helix cincta*, *H. ligata*, *H. figulina*, *H. pomatia* (CZ), and *H. thessalica* (RO)

The mitogenomic sequences were assembled from RNA sequencing reads. RNA was extracted from posterior foot tissues preserved in RNAlater. Libraries were constructed from poly(A) enriched RNA and sequenced on the NovaSeq X platform (150 bp paired-end). After running rCorrector to correct likely erroneous k-mers and removing reads flagged as having an "unfixable_error", the raw reads were trimmed and filtered with fastp (`--adapter_sequence=AGATCGGAAGAGCACACGTCTGAACTCCAGTCA --adapter_sequence_r2=AGATCGGAAGAGCGTCGTGTAGGGAAAGAGTGT --trim_poly_g --`

`cut_front --cut_tail --cut_window_size 4 --cut_mean_quality 20 --qualified_quality_phred 15 --unqualified_percent_limit 40 --n_base_limit 5 --length_required 15`). *De novo* assembly was performed with Trinity and mitochondrial contigs were identified and combined into longer sequences as detailed above with the help of the *H. pomatia* mitogenome sequence as a reference. To check the quality of the resulting assembly we mapped reads with Bowtie2 and then inspected the mapped visually for problems as above.

*Caucasotachea atrolabiata*, *Lindholmia nordmanni*, *Levantina escheriana*, *Maltzanella dickhauti*, *Helix godetiana*, *H. salomonica*, *H. nucula*, *H. pronuba*, *H. albescens*, *H. philibinensis* and *H. buchii*

Sequencing libraries were prepared from total DNA with the NEBNext® Ultra™ II FS DNA Library Prep Kit for Illumina and sequenced on the NextSeq2000 platform (paired-end, 150 bp). Trimming and filtering was performed as described for *H. pathetica* and *H. pomacella*. Mitogenome assemblies were performed with MITObim. As reference, we provided a chimeric sequence consisting of *H. pomatia* mitogenome with partial *rrnS* and *cox1* replaced with sequences of the respective species. MITObim was run with `--kbait 31`. However, this general strategy was not sufficient in all cases. For *H. godetiana*, the sample with the worst quality of input DNA and consequently the lowest number of reads, we used `--kbait 21` and a reference that also contained *rrnL* from *H. godetiana*. The resulting assembly was missing a large stretch around *nd2* and *nd4*, which was closed by running MITObim again with --kbait 18 and a new reference made as a chimaera of the previous assembly and *H. pomatia* sequence. In *L. nordmanni*, *nd4L* and parts of adjacent genes were missing from the first assembly, so an analogous strategy was used to close the gap here as well as similar gaps in other sequences. In both *H. pronuba* and *H. salomonica*, MITObim never got through iteration 2 in a reasonable time, so we produced a chimaera of the assembly from the first iteration (after deletion of probably non-homologous part towards the 3' end) and used that as reference. In *H. pronuba*, this was repeated several times. Some of the gaps were only apparent from alignments with other species, as were also two cases of misassembly of protein-coding sequences. Re-running MITObim while providing as reference the previous assembly with the missing or misaligned part replaced with Ns helped in most cases. The only problematic issue was found in *H. pronuba*, in which the original assembly of *cox3* contained frameshift deletions. This was eventually resolved by building a BLAST database from the forward reads and using tblastn to search for *cox3* reads with an amino-acid sequence from *H. cincta* as a query. The reads found were then used to assemble the gene manually. There is apparently a duplication of *cox3* in *H. pronuba*, as was supported by the roughly two times higher coverage for *cox3* than the rest of the mitogenome. We assembled the second copy and the flanking regions by manually assembling a part of it from the mapped reads and then running MITObim with this short assembly (877 bp) as a reference and `--kbait 100`.

Camacho C, Coulouris G, Avagyan V, *et al*. BLAST+: Architecture and applications. *BMC*

*Bioinformatics* 2009; ***10***: 421. https://doi.org/10.1186/1471-2105-10-421
Chen S, Zhou Y, Chen Y, *et al*. fastp: An ultra-fast all-in-one FASTQ preprocessor. *Bioinformatics* 2018; ***34***: i884–i890. https://doi.org/10.1093/bioinformatics/bty560
Grabherr MG, Haas BJ, Yassour M, *et al*. Full-length transcriptome assembly from RNA-Seq data without a reference genome. *Nature Biotechnology* 2011; ***29***: 644–652. https://doi.org/10.1038/nbt.1883
Groenenberg DSJ, Duijm E. The complete mitogenome of the Roman snail *Helix pomatia* Linnaeus 1758 (Stylommatophora: Helicidae). *Mitochondrial DNA Part B* 2019; ***4***: 1494–1495. https://doi.org/10.1080/23802359.2019.1601512
Kearse M, Moir R, Wilson A, *et al*. Geneious Basic: An integrated and extendable desktop software platform for the organization and analysis of sequence data. *Bioinformatics* 2012; ***28***: 1647–1649. https://doi.org/10.1093/bioinformatics/bts199
Korábek O, Petrusek A, Rovatsos M. The complete mitogenome of *Helix pomatia* and the basal phylogeny of Helicinae (Gastropoda, Stylommatophora, Helicidae). *ZooKeys* 2019; ***827***: 19–30. https://doi.org/10.3897/zookeys.827.33057
Krueger F. TrimGalore, version 0.6.10. 2020. https://github.com/FelixKrueger/TrimGalore (15 February 2025, date last accessed)
Langmead B, Salzberg SL. Fast gapped-read alignment with Bowtie 2. *Nature Methods* 2012; ***9***: 357–359. https://doi.org/10.1038/nmeth.1923
Li H, Handsaker B, Wysoker A, *et al*. The Sequence Alignment/Map format and SAMtools. *Bioinformatics* 2009; ***25***: 2078–2079. https://doi.org/10.1093/bioinformatics/btp352
Martin M. Cutadapt removes adapter sequences from high-throughput sequencing reads. *EMBnet.Journal* 2011; ***17***: 10. https://doi.org/10.14806/ej.17.1.200
Okonechnikov K, Golosova O, Fursov M. Unipro UGENE: A unified bioinformatics toolkit. *Bioinformatics* 2012; ***28***: 1166–1167. https://doi.org/10.1093/bioinformatics/bts091
Scheel BM, Hausdorf B. Survival and differentiation of subspecies of the land snail *Charpentieria itala* in mountain refuges in the Southern Alps. *Molecular Ecology* 2012; ***21***: 3794–3808. https://doi.org/10.1111/j.1365-294X.2012.05649.x
Sokolov EP. An improved method for DNA isolation from mucopolysaccharide-rich molluscan tissues. *Journal of Molluscan Studies* 2000; ***66***: 573–575. https://doi.org/10.1093/mollus/66.4.573
Song L, Florea L. Rcorrector: Efficient and accurate error correction for Illumina RNA-seq reads. *GigaScience* 2015; ***4***: 48. https://doi.org/10.1186/s13742-015-0089-y
