## Appendix 2 for "Finding the relatives of a critically endangered island endemic reveals a heterogeneous evolution of land snail mitogenomes (Gastropoda: Helicidae)"

SUPPLEMENTARY APPENDIX S2

### Genital system anatomy of *Isaurica callirhoe* and *I. lycia*

The three *Isaurica* species whose anatomies are known, *I. lycia*, *I. pamphylica* and *I. schuetti*, cannot be separated by the gross anatomies of their genitalia (Pfeiffer & Wächtler, 1939; Subai, 1994). Subai (1994) mentioned only minor differences among the individual organs of the species. For example, the mucus glands of *I. lycia* had fewer (2–3) branches than those of *I. pamphylica* (3–5) but intraspecific (and intraindividual) variation renders this an unreliable trait (see the drawings of Subai, 1994 and Pfeiffer & Wächtler, 1939). Subai (1994) considered the shape of the atrial stimulator (“Penis-Reizkörper”), located in the penis near its opening into the atrium, to be a valuable character. It should be spherical in *I. lycia* and flat in *I. schuetti* and *I. pamphylica*. But such descriptions can be subjective or affected by preservation and it is difficult to determine the shapes of the stimulators reliably from Subai's drawings.

We compared the genital system anatomy of our sample of *I. callirhoe* with *I. lycia* from Megisti (Kastellorizo) to rule out any substantial differences that might suggest that *I. callirhoe* is a separate species. Unfortunately, when the snail was drowned for preservation, it died with its atrium everted, and the bursa apparatus has been lost during dissection. The length of the penis could thus be only approximately measured due to the eversion of the atrium and the shape of the atrial stimulator could not be observed. There was no dart in the dart sac. One mucus gland had two branches (the second one was damaged during dissection).

Despite being only partially able to examine the genitalia of *I. lycia*, we conclude that there are likely no significant differences and the genital system anatomy is consistent with that of other *Isaurica* species. The flagellum length in the *I. callirhoe* specimen is similar to the 26 mm given for *I. lycia* from Megisti by Pfeiffer & Wächtler (1939). The observations are summarized in the table and figure below.

The animal was dark, nearly black at the back with a paler, yellow-gray sole of the foot. This is within the range described by Subai (1994) for *I. lycia*: “whitish-beige, or light grey-brown to dark grey, darker coloured on the back and between the tentacles”.

| **measurement (in mm)** | ***Isaurica lycia* (Megisti)** | ***Isaurica callirhoe* (Akdağ)** |
|---|---|---|
| penis | 9.9 | ? |
| penis + distal epiphallus | 15.2 | >10 |
| proximal epiphallus | 5.7 | 6.2 |
| flagellum | 19.7 | 28.6 |
| vagina | 9.7 | 5 |
| free oviduct | 4.2 | 2 |
| diverticulum | 50.0 | lost |
| proximal pedunculus of bursa | 40.4 | lost |
| distal pedunculus of bursa | 10.2 | lost |

on the next page: *Isaurica callirhoe* from Akdağ, collected 6.5.2007 (A–C) and *Isaurica lycia* from Megisti, collected 2.12.1996 near the village (D, E) A, head of a preserved *I. callirhoe* specimen with the atrium everted; B, live individual of *I. callirhoe*; C, the internal anatomy of penis of *I. callirhoe*; D, the internal anatomy of penis of *I. lycia*; E, the genital system of *I. lycia* showing the measured organs. Abbreviations: dpd = distal pedunculus; ds = dart sac; dv = diverticulum; ep = epiphallus; fl = flagellum; mg = mucous glands; pn + dep = penis + distal epiphallus, pep = proximal epiphallus; pp1 = proximal penial papilla; pp2 = distal penial papilla; ppd = proximal pedunculus; pr = penial retractor muscle; vd = vas deferens.

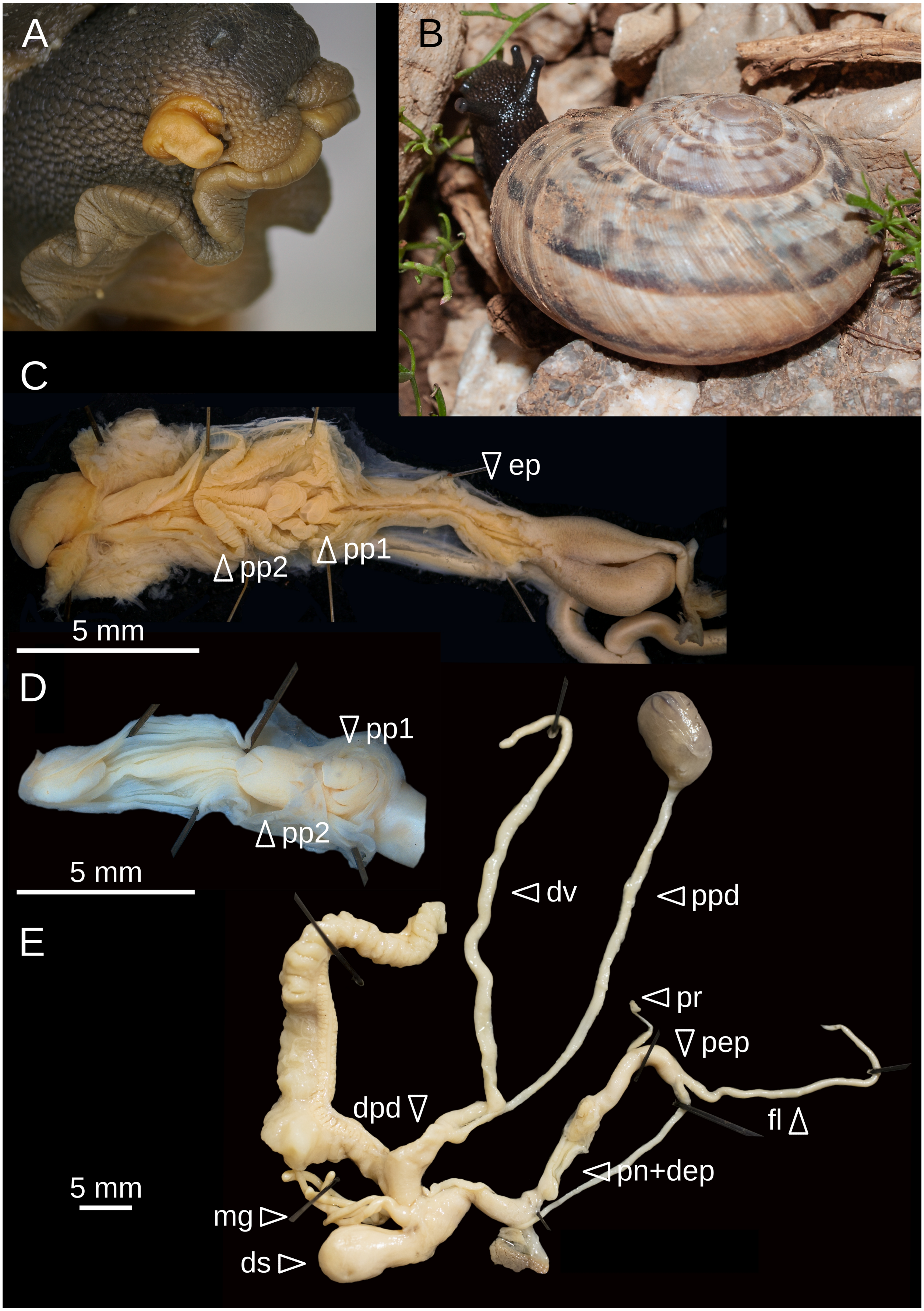
A
B
C
∇ ep
∆ pp2
∆ pp1
5 mm
D
∇ pp1
∆ pp2
5 mm
◁ dv
◁ ppd
E
◁ pr
∇ pep
dpd ∇
fl ∆
◁ pn+dep
5 mm
mg ▷
ds ▷
